# Parallel measurement of transcriptomes and proteomes from same single cells using nanodroplet splitting

**DOI:** 10.1101/2022.05.17.492137

**Authors:** James M. Fulcher, Lye Meng Markillie, Hugh D. Mitchell, Sarah M. Williams, Kristin M. Engbrecht, David J. Degnan, Lisa M. Bramer, Ronald J. Moore, William B. Chrisler, Joshua Cantlon-Bruce, Johannes W. Bagnoli, Wei-Jun Qian, Anjali Seth, Ljiljana Paša-Tolić, Ying Zhu

## Abstract

Single-cell multiomics provides comprehensive insights into gene regulatory networks, cellular diversity, and temporal dynamics. Here, we introduce nanoSPLITS (nanodroplet SPlitting for Linked-multimodal Investigations of Trace Samples), an integrated platform that enables global profiling of the transcriptome and proteome from same single cells using RNA sequencing and mass spectrometry-based proteomics, respectively. Benchmarking of nanoSPLITS demonstrated excellent measurement precision, with deep proteomic and transcriptomic profiling of single-cells. We applied nanoSPLITS to cyclin-dependent kinase 1 inhibited cells and found phospho-signaling events could be quantified alongside global protein and mRNA measurements, providing new insights into cell cycle regulation. We also extended nanoSPLITS to single-cells isolated from human pancreatic islets, introducing an efficient approach for facile identification of unknown cell types, and detecting their protein markers by mapping transcriptomic data to existing large-scale single-cell RNA sequencing reference databases. Herein, we establish nanoSPLITS as a new multiomic technology incorporating global proteomics and anticipate the approach will be critical to furthering our understanding of single-cell systems.

## Introduction

Multicellular organisms contain a variety of cell populations and subpopulations, which are well-organized in defined patterns to implement critical biological functions. The development and rapid dissemination of single-cell omic technologies have dramatically advanced our knowledge on cellular heterogeneity,^1-3^ cell lineages,^4^ and rare cell types.^5^ However, most existing technologies only capture single modalities of molecular information. Such measurement provides only a partial picture of a cell’s phenotype, which is determined by the interplay between the genome, epigenome, transcriptome, proteome, and metabolome. Moreover, proteins are of particular interest in establishing cellular identities because they are the downstream effectors and their abundance cannot be reliably inferred from other modalities, including transcript^6^. Unfortunately, existing multimodal transcriptome-proteome measurements ^7-14^ are restricted to at most a few hundred (and often significantly fewer) protein targets. These measurements also require intermediate antibodies to recognize epitopes, which can be limited by availability and specificity^15,16^.

A route for overcoming these limitations is through the adoption of a mass spectrometry (MS)-based approach. With the advance of microfluidic sample preparation^17^ and isobaric labeling^18^, single-cell proteomics (scProteomics) is now capable of measuring thousands of proteins from a single cell.^19-21^ Encouraged by these developments, we sought to acquire multimodal transcriptome-proteome measurements from the same single cells by integrating single-cell RNA sequencing (scRNAseq) with scProteomics. To enable efficient integration, we developed nanoSPLITS (nanodroplet SPlitting for Linked-multimodal Investigations of Trace Samples), a method capable of dividing single-cell lysates via two nanoliter droplet microarrays and separately measuring them with scRNAseq and MS-based scProteomics. NanoSPLITS builds on the nanoPOTS platform that allows for high-efficiency proteomic preparation of single-cells by miniaturizing the assay to nanoliter scale volumes^19,22^. We have previously demonstrated reaction miniaturization not only reduces non-specific adsorption-related sample losses, but also enhances enzymatic digestion kinetics.^22^ Similarly, we reason the use of nanoliter droplets can improve overall sample recovery of both mRNA transcripts and proteins for sensitive and comprehensive single-cell multiomics.

Herein, we demonstrated that nanoSPLITS-based multiomics can provide comprehensive molecular characterizations of both secondary and primary cells with insights including covarying functional protein/gene clusters and unique phosphorylation events, which are usually unavailable in a single-omics measurements. The capability to obtain the proteome profiles of individual cell types in complex tissue dissociations is highly desirable for therapeutic target discovery and biomarker identification. However, it is highly challenging to measure many single cells with label free LC-MS-based proteomics workflows. To alleviate this problem, we developed an efficient approach for cell-type annotation and protein marker detection by mapping nanoSPLITS transcriptomics data to existing scRNAseq reference databases. This reference mapping approach significantly reduces the required numbers of cells for clustering-based cell-type annotation.

## Results

### Optimization of nanoSPLITS provides deep transcriptome and proteome coverage in single cells

The overall workflow of the nanoSPLITS-based single-cell multiomics platform is illustrated in **Fig. 1**. Briefly, we employed an image-based single-cell isolation system to directly sort single cells (contained in ∼400 pL volume droplets) into lysis buffer (200 nL), followed by a freeze-thaw cycle to achieve cell lysis. Notably, the selective hydrophobic/hydrophilic patterning of nanoSPLITS chips ensures that droplets are constrained to the 1.2 mm wells on the chip. Next, the microchip containing single-cell lysate is manually aligned with a separate chip containing only cell lysis buffer. The droplet arrays on the two chips are merged for 15 seconds and then separated, before repeating twice (**<u>Movie S1</u>**). As nanoSPLITS chip contains 48 wells of 1.2 mm diameter separated by 4.5 mm of space, chips can easily be aligned to accomplish droplet merging. The chip receiving lysate ( “acceptor” chip) was then transferred into a 384-well plate for scRNAseq with Smart-seq 2 protocol^1^, while the chip initially containing the lysate (“donor” chip) is digested using an n-dodecyl-β-D-maltoside (DDM)-based sample preparation protocol and directly analyzed with an ion-mobility-enhanced MS data acquisition method for scProteomics^23^.

**Fig.1:**
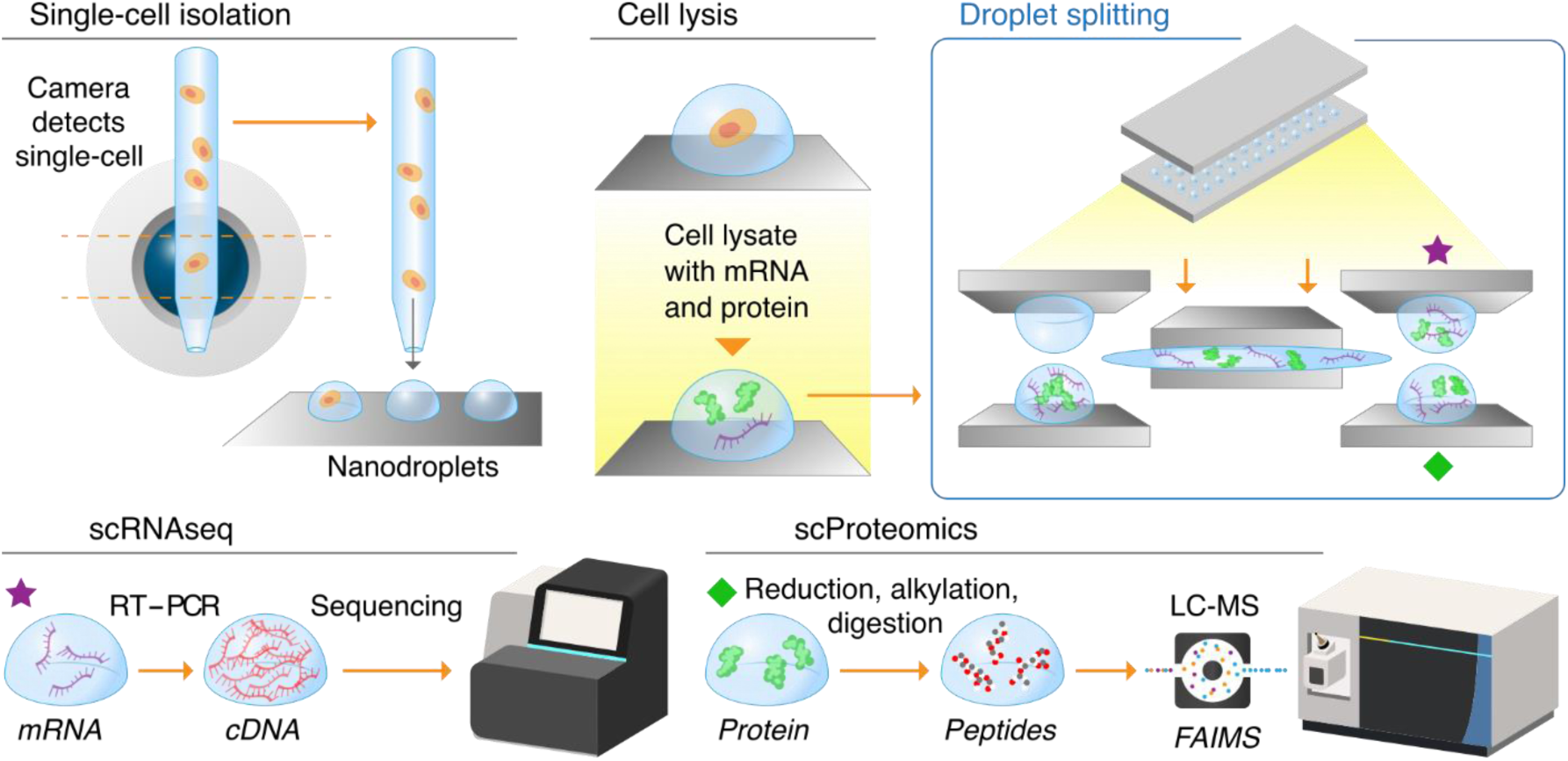
Overview of the nanoSPLITS-based single-cell multiomics platform. Schematic illustration showing the workflow including cell sorting, lysis, droplet merging/mixing, and droplet separation for downstream scRNAseq and scProteomics measurement.

We first optimized cell lysis buffer conditions to ensure compatibility with both proteomic and transcriptomic workflows. Typically, scProteomics utilizes a buffer containing DDM to reduce non-specific binding of proteins to surfaces while scRNAseq often includes recombinant protein-based RNase inhibitors to reduce mRNA degradation. To evaluate their impacts on both methods, we tested the inclusion of these additives in a moderately buffered hypotonic solution (10 mM Tris, pH 8) with 20 mouse alveolar epithelial cells (C10). The inclusion of 1 x RNase inhibitor modestly suppressed proteomic identifications, while 0.1% DDM had no significant impact on transcript identifications (**<u>Fig. S1a</u>** and **<u>S1b</u>**). Furthermore, removal of RNase inhibitor from RNAseq analysis had minimal effect on gene identifications (**<u>Fig. S1a</u>**). Therefore, we decided to use 0.1% DDM (w/v) in 10 mM Tris solution as the cell lysis buffer for our initial nanoSPLITS experiments.

Next, we evaluated the split ratios between two 200-nL droplet arrays. Using fluorescein as an initial model, the nanoSPLITS procedure achieved splitting ratios between 46% to 47%, with 50% representing a perfectly equal split (**<u>Fig. S2</u>** and **<u>Table S1</u>**). As the properties of fluorescein may not fully model proteins, we split 10 C10 cells (n = 6) and quantified protein abundances in both chips with LC-MS. Encouragingly, we found the split ratios to be consistent and precise across the chips, with a median CV of 0.12 for the 10 pooled cell samples (**<u>Fig. S3a</u>**). Using the mean protein abundance for each protein identified from both chips, we determined the relative proportion of each protein found on either chip (**<u>Fig. S3b</u>**). Surprisingly, the median proportion of protein retained on the donor chip was ∼75%, greater than what would be expected in a perfect split. One potential explanation is that the cell lysate may require more time for diffusion between the merged droplets, although the diffusion coefficient of GFP in water (∼100 µm^2^s^-1^) suggests merging on the order of seconds should be sufficient. In addition, a diffusion-based effect would be expected to have a size dependence, which is not apparent based on the proteins quantified (**<u>Fig. S3c</u>**). Another explanation might be that the modest hydrophobic coating of the nanowells captured proteins on the surface, resulting in more proteins retained on the donor chip. Thus, we took advantage of this outcome and chose the donor chip for scProteomic measurements to obtain deeper proteome coverages (**Fig. 1**).

We next sorted several quantities of C10 cells (groups of 11, 3, and 1 cell into separate wells) and measured them using the multiomics workflow. Considering a minimum of 5 reads per gene for transcript identifications and 1% FDR cutoff for protein identifications (5% at ion level during match-between runs, MBR), robust coverages of genes and proteins were achieved across all tested conditions (**Fig. 2a**). As expected, coverage reduced with decreasing cell numbers. Single-cell transcriptome and proteome measurements provided 5,848 and 2,934 identifications on average, respectively. To ensure nanoSPLITS did not introduce bias toward different cellular components due to the nanodroplet splitting process, we investigated the distribution of gene and protein identifications from single cells within cellular component ontology (CCO). We found scProteomics and scRNAseq had corresponding identifications within cellular components that encompassed all major organelles (**Fig. 2b**). Furthermore, 25% of the proteins from the scProteomics analyses had CCO localizations to the nucleus, 219 of which have known roles in transcription. This is notable as nuclear proteins are typically drivers in gene regulation and transcription, and current multimodal technologies have been limited in the ability to directly measure nuclear protein abundances.

**Fig. 2:**
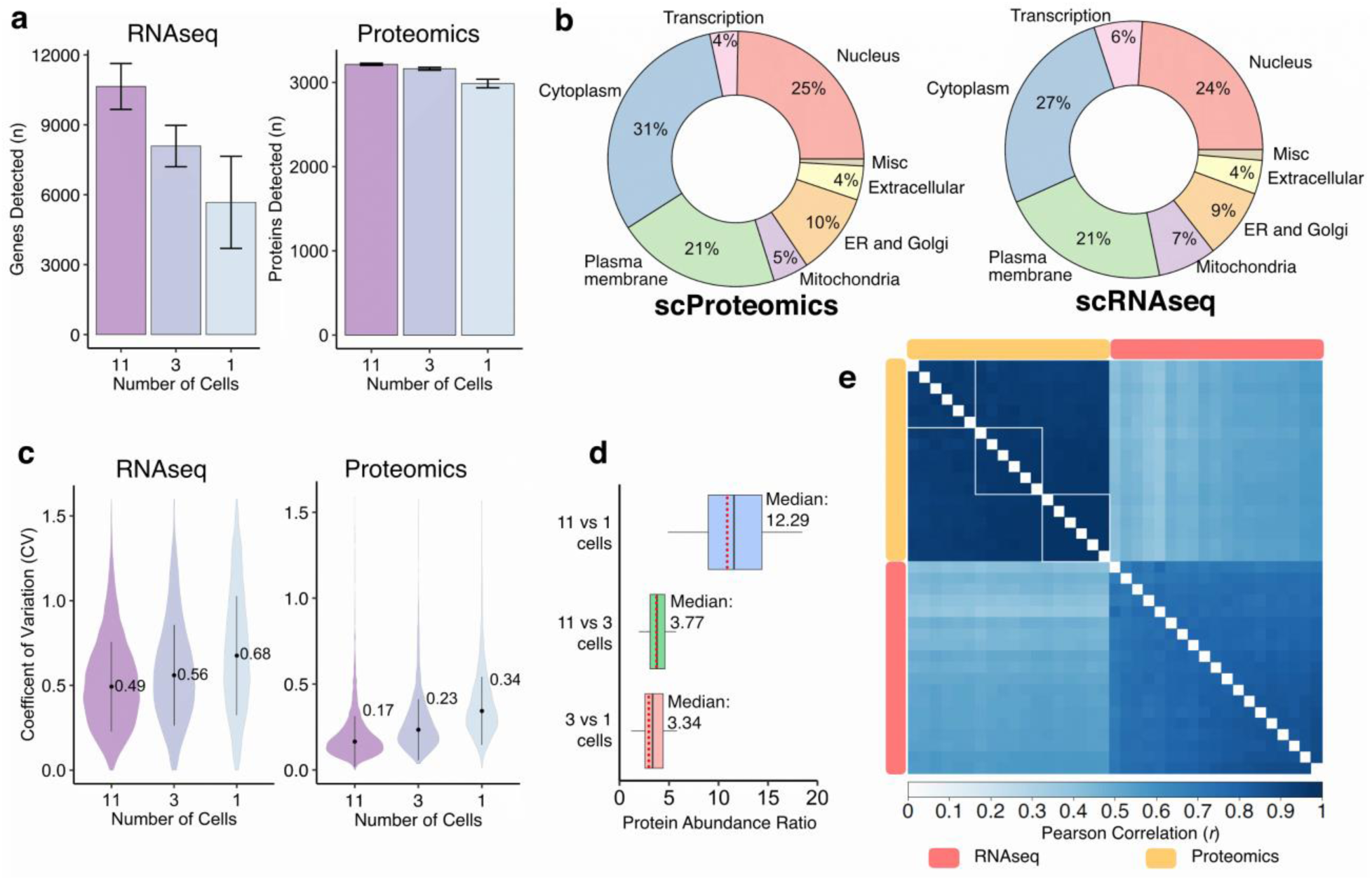
Qualitative and quantitative assessment of nanoSPLITS for transcriptome and proteome measurements. (**a**) Mean number of detected genes and proteins for each modality (n = 6, 6, and 7 for 11, 3, and 1 C10 cell, respectively). Error bars indicate standard deviations (±s.d.).(**b**) CCO distributions of genes (scRNAseq) and proteins (scProteomics) identified in the single-cell data (n = 7). (**c**) Distributions of the coefficients of variation (CV) for all genes and proteins with at least 2 observations across replicates. Indicated values represent median CVs, which are also indicated at the center point within each distribution. (**d**) The ratios of protein abundance were calculated for comparisons between the different pooled cell samples (11 vs 1, 11 vs 3, and 3 vs 1). Experimental medians are indicated at the black crossbar while the theoretical ratio for each comparison is shown at the red dotted line within each boxplot. (**e**) Pearson correlation heatmap with clustering of transcriptomics and proteomics results. Only the proteomic data showed complete clustering of 11,3 and 1 cell samples, indicated by the inscribed white squares. Self-comparisons along the diagonal are excluded (white).

We next evaluated the quantitative reproducibility for each modality by calculating the coefficients of variation (CVs) of transcriptome and proteome abundances. Median transcriptome CVs ranged from 0.49 for 11 cells to 0.68 for single cells, while proteome median CVs ranged from 0.17 for 11 cells to 0.34 for single cells (**Fig. 2c**). The trend of increasing CVs from 11 to 1 cell was expected, as the mixed cell populations represent averages of underlying cell to cell variation. Notably, the median CV (0.68) for single-cells measured by Smart-seq2 were higher than the corresponding scProteomic median CV (0.34). This is also in agreement with recent reports that have shown median CVs across single-cells to be as high as 1.08 ^19,21,24^. These higher CVs partially reflect the dynamic nature of mRNA transcript expression relative to their protein counterparts, which have longer half-lives on average, high copy numbers, and therefore can be more precisely measured ^25^. We also compared the ratios of the measured protein abundances between the different cell populations. Encouragingly, the experimental fold differences between the median intensities for 11, 3, and 1 C10 cell are very close to the expected theoretical values (**Fig. 2d**). For example, the median protein abundance ratio for 3 cells compared to single-cells was 3.34, within 12% of the theoretical 3-fold difference. These results provide solid evidence that nanoSPLITS-based single-cell multiomics platform can provide sensitive and reproducible quantitative measurements of both the transcriptome and proteome of the same single cells.

Finally, we determined the Pearson correlation coefficients (*r*) across and within modalities using conceptually similar normalized transformations for each modality (**Fig. 2e** and **<u>Fig. S4</u>**; TPM, transcripts per million for transcriptomics, and riBAQ, relative intensity-based absolute quantification for proteomics). In line with the CV distributions (**Fig. 2c**), proteomics data had better agreement between samples compared with transcriptomics data, once again highlighting the relatively stable nature of the proteome and the dynamic nature of the transcriptome where many genes are often expressed in short transcriptional “bursts” ^25^. The cross-correlation between the transcriptome and proteome in single cells was moderate with most correlation coefficients falling within the range of 0.35 to 0.45, on par with previous reports ^19,21,25^.

### Classification of cell types and identification of markers with nanoSPLITS

Having established baseline characteristics of multimodal data, we then applied nanoSPLITS to analyze two cell lines: mouse epithelial (C10) and endothelial cells (SVEC). As nanoSPLITS uses only a portion of the total cellular content, we sought to ensure our multimodal measurements could distinguish cell types and detect gene/protein markers.

As shown in **Fig. 2a**, both cell types and modalities were easily clustered based on correlations alone. Within-modality correlations were higher in scProteomics than scRNAseq for both cell types (**<u>Fig. S5a</u>**). Cross-modality correlation analysis produced a slightly broader range of *r* (**<u>Fig. S5a</u>** and **<u>Fig. S6a</u>**). We also compared the cross-modality correlations between same single-cells (“intracell”) and different single-cells (“intercell”), however there appeared to be no significant difference between them (**<u>Fig. S5a</u>**). Overall, SVEC cells exhibited lower correlations which we attribute to their smaller cell size and correspondingly reduced measurement depth/precision as it is known that protein content is linked to cell size (**<u>Fig. S5b</u>** and **<u>Fig. S6a</u>**)^26^. The protein/gene overlap analysis clearly shows how measurement depth is strongly linked to cell size (**<u>Fig. S5b</u>**). On average, C10 cells had ∼1,800 overlapping identifications while SVEC cells had ∼900 overlapping identifications between scRNAseq and scProteomic modalities. All proteins (3,609) detected in the scProteomic data could be detected as mRNA transcripts in the scRNAseq data; and, as expected, the majority of detected proteins were derived from mRNA transcripts of higher abundance (**<u>Fig. S6b</u>**). Finally, we calculated mRNA-protein correlations for each gene that was observed in both modalities. The distribution of mRNA-protein correlations was statistically different compared to a distribution of randomly sampled correlations, albeit modestly (**Fig. 2b**).

We also evaluated if the multiomics data could be used to identify cell-type-specific marker genes and proteins. **Fig. 2c** shows the top 5 significantly enriched genes and proteins for each cell type. Interestingly, the overlap between scProteomics and scRNAseq of these significant markers was relatively low, indicating the widely used scRNAseq method may not be sufficient to provide reliable marker genes for protein-based functional assays. Encouragingly, the previously established SVEC-cell marker *H2-K1* was identified at both the protein and mRNA level (**Fig. 2c**). Dimensionality reduction with principal component analysis (PCA) delineated both cell types for scRNAseq and scProteomics despite the cell contents being divided (**<u>Fig. S7</u>**). The integration of two modalities through a weighted nearest neighbor (WNN)^27^ analysis provided robust clustering in the two-dimensional space and visual confirmation of SVEC marker H2-K1 (**Fig. 2d**). Using canonical cell cycle markers,^28^ we could also identify individual C10 cells in different cell cycle phases, suggesting subtle cell to cell variation was retained after the droplet splitting process (**Fig. 2e** and **<u>Fig. S8</u>**). For example, cyclin-dependent kinase 1 (*Cdk1*) is upregulated at the transcriptional level in S and G2M phase C10 cells (**Fig. 2e**). Several other established cell cycle phase genes demonstrated similar differential abundance at the transcriptional level, including DNA topoisomerase IIα (*Top2a*), cyclin B1 (*Ccnb1*), G2 and S phase expressed protein 1 (*Gtse1*), cytoskeleton-associated protein 5 (*Ckap5*), and anillin (*Anln*) (**<u>Fig. S8</u>**). Furthermore, Top2a protein appeared to be increased in G2/M cells as well.

### Characterization of cell cycle features through the integration of scRNAseq, scProteomics, and scPhosphoproteomics

Next, we investigated whether nanoSPLITS could be applied to study biological systems with more subtle differences. To accomplish this, we characterized cell-cycle features of C10 cells by arresting them in G2/M phase with a CDK1 inhibitor (RO-3306) and comparing them to an untreated C10 cell population^29^. NanoSPLITS facilitated comprehensive proteome and transcriptome measurements, with an average of 2,942 proteins and 5,559 genes identified in G2/M arrested cells as well as 2,574 proteins and 4,173 genes for untreated cells. The higher number of identifications for G2/M arrested cells can be attributed to their larger size (and therefore higher protein/mRNA content), which was noted during cell sorting (**<u>Fig. S9</u>**). Although no phosphopeptide enrichment was performed^30^, the FragPipe proteomics pipeline with FDR-controlled MBR also identified over 300 unique phosphopeptides, of which 138 were reproducibly observed with <50% missing values. G2/M arrested and untreated cells could easily be clustered and separated by dimensional reduction (PCA and WNN-UMAP) for scProteomic and scRNAseq data (**<u>Fig. S10</u>**). Overall, 3,182 proteins and 4,186 mRNA transcripts were tested for differential abundance, and 1,930 protein/genes had paired data from nanoSPLITS. Quantitative analysis afforded 327 proteins, 1,434 genes, and 29 phosphopeptides that were differentially abundant (log_2_FC > 0.5 or < -0.5 and FDR < 0.01). Importantly, covariant protein and mRNA clusters were identified by hierarchical clustering (**Fig. 4a**, **Fig. 4b**, and **<u>Fig. S11</u>**).

**Fig.3:**
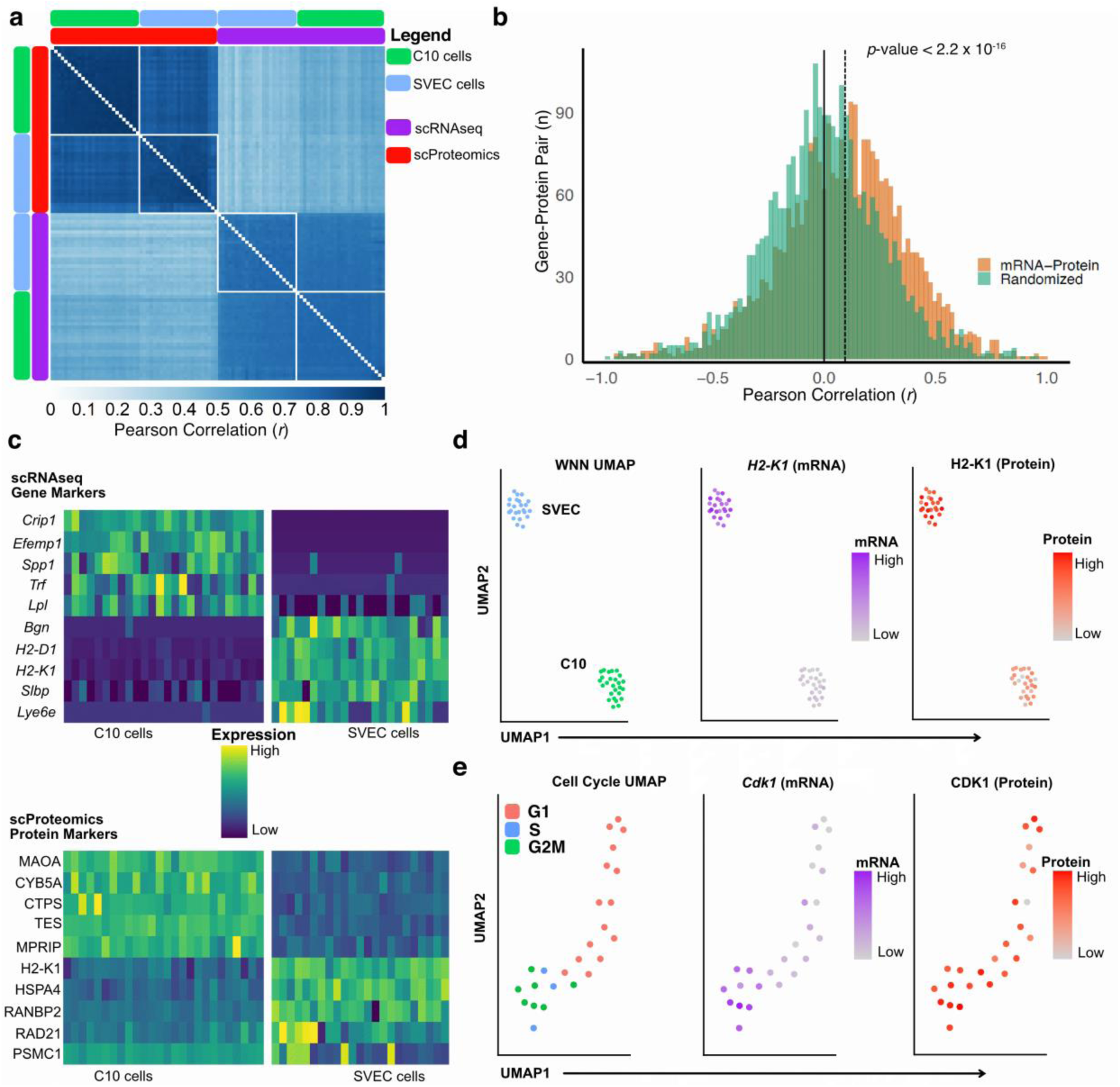
Underlying cell phenotype signatures are maintained after nanoSPLITS. (**a**). Pearson correlation heatmap with clustering of scRNAseq and scProteomic results for both single C10 (n = 26, paired) and SVEC (n = 23, paired) cells. (**b**) Histogram of mRNA-protein correlations for each gene quantified in both modalities with at least 4 observations (performed separately for C10 and SVEC cell types). Statistical testing was performed by non-parametric Mann-Whitney test. Solid and dashed line indicates median of randomized and experimental correlations, respectively. (**c**) Top 5 gene markers from scRNAseq data and protein markers from scProteomics data for each cell type. Candidate marker features were determined using a Wilcoxon Rank Sum test (FDR corrected p-values <0.001). (**d**) Weighted-nearest neighbor (WNN) Uniform Manifold Approximation and Projection (UMAP) generated using Seurat to integrate scRNAseq and scProteomic data. Middle and right panels are colored based on H2-K1 gene (purple) and protein (red) abundance, respectively. (**e**) Feature-based UMAP generated for C10 cells using cell-cycle markers derived from scRNAseq data. Middle and right panels are colored based on Cdk1 gene (purple) and protein (red) abundances, respectively. All abundance values shown in **c**, **d**, and **e** are Z-scores after scaling and centering of data.

**Fig.4:**
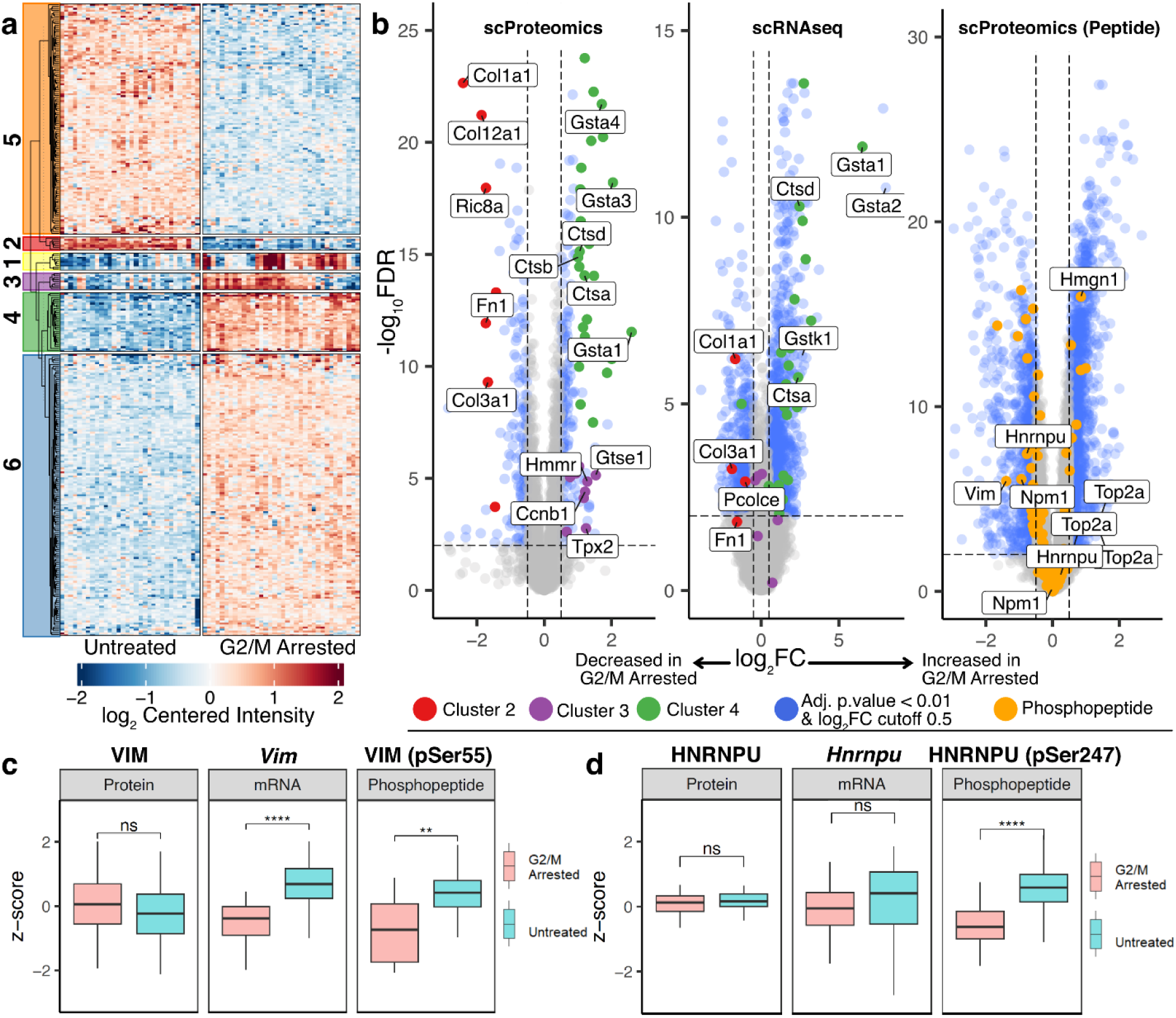
Protein, mRNA, and phosphopeptide cell cycle features from same single cells. **(a)** Clustergram of log_2_ centered intensities for differentially abundant proteins from scProteomic data with FDR < 0.01 and log_2_FC of +/-0.5 (327 proteins). Columns (cells) are clustered by K-means (k = 2), while rows cluster proteins (k = 6). Colored areas along the y-axis indicate protein clusters. (**b**) Volcano plot of G2/M arrested cells/untreated cells for scProteomic data (3,182 proteins quantified), scRNAseq data (4,186 genes with < 50% missing values), and peptide level (16,938 peptides).(**c**) Integration of scRNAseq, scProteomic, and phosphoproteomic data by comparing relative abundances from each modality for VIM. **** FDR < 0.00001, *** FDR < 0.0001, ** FDR < 0.001, and ns indicates not significant. (**d**) Same as (**c**) but for HNRNPU.

Of the proteomic clusters identified, clusters 2, 3, and 4 contained members with the strongest functional relationships. Cluster 2 (**Fig. 4a** and **Fig. 4b**) contained 7 proteins that were significantly more abundant in the untreated C10 cells. These were mainly extracellular matrix (ECM) proteins, including three collagens (COL1A1, COL3A1, and COL12A1), fibronectin (FN1), and procollagen C-endopeptidase enhancer 1 (PCOLCE). The remaining two proteins were endoplasmic reticulum junction formation protein lunapark (LNPK) and Synembryn-A (RIC8A), both of which play important roles within mitosis^31,32^. Cluster 3 provided 13 proteins that were more abundant in G2/M arrested C10 cells, all of which were enriched for mitotic-relevant annotations (**<u>Fig. S12</u>**). Interestingly, cluster 3 did not show concordance between the two modalities which we attribute to the temporal difference between peak translation and protein abundance in preparation for and during mitosis (**Fig. 4b**)^33^. Cluster 4 contained genes with the largest log_2_FC differences at both the mRNA and protein level and was largely composed of glutathione S-transferases and cathepsins (**<u>Fig. S13</u>**). For cluster 2 and cluster 4, strong concordance between the scProteomic and scRNAseq data was noted (**Fig. 4b**, see **Supplementary Information** for further discussion on covariant clusters identified in this study).

From the over 300 phosphopeptides from scProteomic data, 29 were dysregulated in G2/M arrested cells (**Fig. 4b**). We also compared the relative abundance of these phosphopeptides to their corresponding protein and mRNA abundances. Notably, vimentin (VIM) pSer55, a known phosphosite generated by CDK1 typically observed during the transition from prometaphase to metaphase^34,35^, was decreased in abundance significantly in G2/M arrested cells (**Fig. 4c**). This decrease in protein phosphorylation appeared to be independent of the global protein abundance (**Fig. 4c**), providing the most direct evidence for the loss of CDK1 kinase activity inhibited by RO-3306 ^29^. Phopshosite pSer247 on heterogeneous nuclear ribonucleoprotein U (HNRNPU) was also found to be changing independent of overall protein abundance and followed a similar trend as VIM pSer55, suggesting it may be a target of CDK1 (**Fig. 4d**).

### Leveraging large-scale single cell transcriptomics reference databases to confidently annotate cell types and identify functionally relevant protein markers

Having established the precision, depth, and scope of our nanoSPLITS measurements in secondary cell lines, we next applied nanoSPLITS to dissociated human pancreatic islet cells from two donors. The overall identification depth across the dissociated islet cells was comparable to C10 and SVEC cells, with an average of 1,775 genes and 1,949 proteins identified per cell. A notable advantage of having parallel scRNAseq data from nanoSPLITS measurements is the ability to annotate cells via reference maps from published scRNAseq experiments. By integrating nanoSPLITS scRNAseq data into the reference-based mapping pipeline in Seurat (Azimuth)^36,37^, we could assign nine different cell types across 106 cells (**Fig. 5a**). These cell types encompassed both pancreatic exocrine (acinar and ductal cells) and endocrine (alpha, beta, delta, and gamma cells) tissue. Notably, resident immune cells (representing only 0.6% of cells in the Azimuth reference atlas) in the pancreas could also be identified. Established protein markers showed strong agreement with the cell types annotated through Seurat (**<u>Fig. S14</u>**). We next trained a sparse partial least squares discriminant analysis (spls-DA) model on the cell types annotated by Seurat (see Supplementary Methods in **<u>Supplementary Information</u>**). Applying the model to unannotated cells afforded 23 more annotated cells from the scProteomic data, with only 8 remaining “unclassified” (**<u>Fig. S15</u>**). Therefore, our analysis encompassed 106 cells with scRNAseq data, 126 cells with scProteomic data, and 95 cells with both modalities (**<u>Fig. S16</u>**).

**Fig.5.**
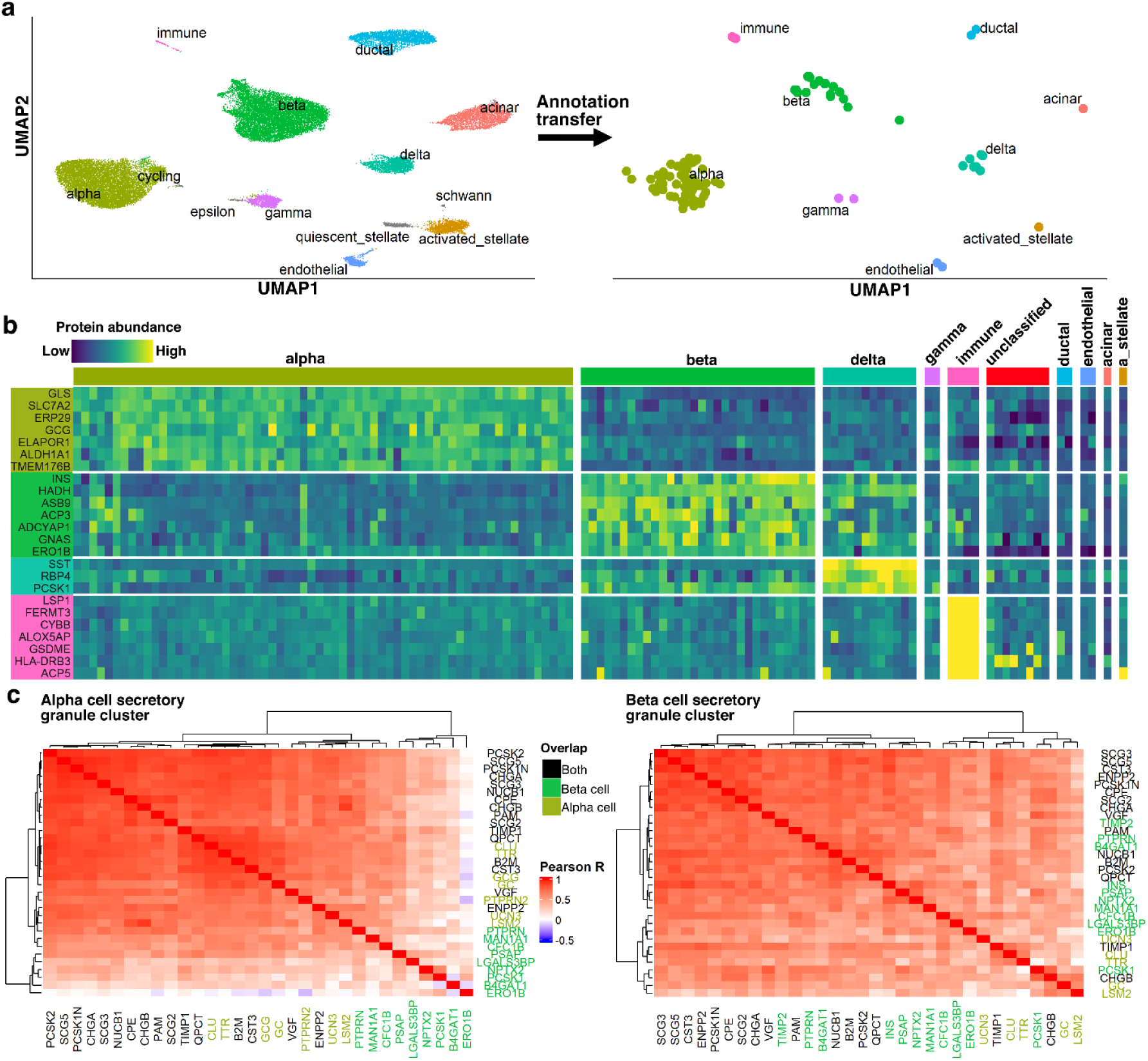
Multiomics analysis of dissociated pancreatic islet cells. **(a)** Left panel: UMAP of Azimuth human pancreatic reference map consisting of 35,289 cells^27^. Cell types are indicated by color and inset text labels. Right panel: UMAP of 106 dissociated pancreatic islet cells from two human donors based on label transfer from Azimuth reference map to nanoSPLITS scRNAseq data. Cells with annotation scores of > 0.8 are shown. (**b**) Abundance heatmap of selected protein markers identified from nanoSPLITS scProteomics data for alpha cells (n = 64), beta cells (n = 30), delta cells (n = 12), immune cells (n = 4), and unclassified cells (n = 8), compared across all annotated cell types. Candidate markers were determined using a Wilcoxon Rank Sum test (adjusted p-values <0.001) (**c**) Clustergram of Pearson correlations for secretory granule proteins hierarchically clustered for alpha (left panel) and beta cells (right panel). “Overlap” indicates if proteins were observed in both alpha and beta cell secretory granule clusters or only one cluster (alpha or beta).

With well-annotated cell types, we then focused on identifying unique protein markers for major endocrine cell populations and immune cells via pairwise abundance comparisons. Many proteins with cell-type specific profiles could be observed (**Fig. 5b**). This includes several of the well-established markers for pancreatic endocrine cells, such as GCG for alpha cells, INS beta cells, and SST for delta cells. Importantly, we also identified several marker proteins that, to the best of our knowledge, have only been established at the mRNA level in single-cells (ALDH1A1, SLC7A2, TMEM176B, and GLS for alpha cells; HADH, ERO1B, and ADCYAP1 for beta cells; RBP4 and PCSK1 for delta cells)^38,39^, as well as proteins that have not been identified as markers in large scale sequencing datasets (ERP29 and ELAPOR1 in alpha cells, ACP3 in beta cells). ASB9 and ALDH1A1 are also noteworthy as they both play a role in beta cell maturation, and GNAS was only recently found to be an important regulator of insulin secretory capacity^40,41^. Owing to the different lineage from which they were derived, immune cells presented with the strongest markers amongst the tested cell types. Like the pancreatic endocrine cells, established mRNA markers were observed (ACP5, HLA-DRB3, ALOX5AP, and LSP1) as well as proteins with more limited mRNA expression (FERMT3 and CYBB)^27^.

Finally, we investigated the functional protein covariation profiles of alpha and beta cells to identify differences in their hormone secretion pathways. Through Gaussian Mixture Model (GMM) clustering of the scProteomic results, a cluster of ∼25 proteins was found in each cell type containing known secretory proteins as well as the major secretory hormones GCG and INS for alpha and beta cells, respectively (**<u>Fig. S17</u>**, and **<u>Fig. S18</u>**). Both clusters were significantly enriched for the gene ontological term “secretory granule” (*adjusted p-value* 2.1 x 10^-6^ for beta cells and 1.6 x 10^-6^ for alpha cells). Proteins from both clusters were then used to construct hierarchical clustergrams specific to alpha cells or beta cells (**Fig. 5c**). From the 33 proteins in these secretory granule clusters, 16 core proteins were found to overlap in both cell types. Proteins uniquely correlated within the alpha cell secretory granule cluster included several of known functional relevance in alpha cells (GC, TTR, and UCN3)^38,42,43^. Of the 11 proteins unique to the beta cell cluster, four were largely uncorrelated in alpha cells (NPTX2, B4GAT1, PCSK1, and ERO1B). NPTX2 has only recently been identified as a beta cell marker at the mRNA level, and ERO1B is an oxidoreductase necessary for proper folding of proinsulin^38,44^. Finally, it is notable that metalloproteinase inhibitor TIMP2 deviates significantly from the related TIMP1, which appears to be a core protein in both secretory pathways. TIMP2 is frequently observed in beta cells (28 out of 30 beta cells) and unique to the beta cell secretory granule cluster, however only sparsely observed in alpha cells at much lower abundance (18 out of 64 alpha cells). The role of TIMP proteins in hormone secretion appears to be relatively unknown, however TIMP1 is at least known to localize to compartments resembling secretory vesicles in neutrophils^45^.

## Discussion

Compared with previous technologies that utilize antibodies to infer protein abundances in multimodal experiments, the nanoSPLITS platform employs MS to globally detect proteins and post-translational modifications (PTMs). We have established how the nanoSPLITS approach can enable multimodal profiling of thousands of transcripts and proteins, as well as hundreds of phosphopeptides from same single cells. Quantification of protein abundances from 11, 3, and 1 C10 cells verified experimental abundance ratios were close to theoretical and highly precise. Despite splitting cellular contents, we found the molecular coverage to be broadly distributed across different cellular compartments for both modalities, and global across-modality correlations were in line with prior studies^25^. Finally, applying nanoSPLITS to both secondary cell lines and human primary cells enabled delineation of different cell types and identification of candidate marker genes and/or proteins.

Although our experimental design was not targeted towards clarifying the relationship between mRNA and protein abundance, we could calculate the correlations for all genes with corresponding transcript-protein measurements derived from same single cells (C10 and SVEC in **Fig. 2b**, as well as dissociated pancreatic islet cells in **<u>Fig. S19</u>**). The mRNA-protein correlations measured could be distinguished from distributions of randomly sampled measurements, and a trend towards more positive correlations was observed. However, the medians of the distributions were only modestly shifted from 0, providing further evidence that the lack of correlation between transcript and protein levels is a general phenomenon^11,46^. Indeed, the inherent stochasticity of gene expression is well documented^47^. Overall, our results agree with the perspective that correlations between transcripts and proteins at the single-cell level are less correlated relative to bulk measurements which “average out” different sources of variance within a population of cells^6,11^. However, modest shifts in correlations towards positive *r* suggests that at least some transcripts are more tightly linked to their translated products. We anticipate that sampling larger sets of cells will provide new insights into regulation of protein expression.

We also demonstrated nanoSPLITS can identify subtle cell phenotypes, such as mitotic cell cycle phases. By comparing cells arrested in G2/M via the CDK1 inhibitor RO-3306 to an actively cycling C10 population, we could identify abundance changes for proteins and transcripts in line with the expected phenotype, as well as covarying protein/gene clusters relevant to mitotic processes. Over 300 phosphopeptides were also identified without phosphopeptide enrichment, providing one of the deepest single-cell phosphoproteomic datasets to date^30,48^. From this dataset, we found phosphosites pSer55 VIM and pSer247 HNRNPU were strongly impacted by CDK1 inhibition. Interestingly, the phosphorylation of HNRNPU by PLK1 at a separate site (pSer59) is known to be a key step in mitosis that requires an upstream priming phosphorylation event by CDK1, but the exact site that CDK1 acts on HNRNPU has yet to be identified^49^. Our data suggests this upstream priming by CDK1 may be positioned at Ser247. These phosphoproteomic observations demonstrate how additional insight can be acquired from PTMs ^30^, and exemplify the benefits of MS-based proteomics in single-cell analysis.

Finally, we developed a powerful approach to annotate unknown cell types by leveraging large-scale scRNAseq reference database with nanoSPLITS multiomic data. This highlights a unique capability of single-cell multimodal analyses: utilizing one modality (e.g., transcripts) as a bridge to inform cell types. This allows for discovery of cellular markers in other modalities (e.g., proteins). Such capability is particularly useful for identifying protein markers in complex cell populations with scProteomics, as only a small number of single cells need to be measured. Additionally, the approach avoids having to use the same data for both clustering and differential expression analysis (a process referred to as “double dipping” that inflates false-positives)^50^. By performing nanoSPLITS on human pancreatic islet cells, we demonstrated how most islet cell types, including rare cell types such as immune cells, could be confidently annotated. We validated previously identified transcript markers and found several novel protein markers, notably including protein ASB9. Our results extend knowledge on ASB9 by showing it is maintained in mature beta cells and not just transiently expressed as part of terminal beta cell maturation^40^. With the ongoing efforts in large-scale single-cell mapping such as Human Cell Atlas and HuBMAP^51,52^, reference maps for many organs are readily available. Thus, we envision the integration of reference mapping with nanoSPLITS multiomics as a powerful approach for discovery proteomics in heterogeneous cell populations.

The nanoSPLITS platform holds promise to become a powerful discovery tool for biomedical applications, facilitating characterization of heterogeneity in tissues, peripheral blood cells, and circulating tumor cells. Although a low throughput approach was employed in this study, high-throughput multiplexing approaches such as CEL-Seq^53^ for transcriptomics and SCoPE-MS^18^ for proteomics can
 be readily integrated into the nanoSPLITS workflow. The technique is also not restricted to the two modalities demonstrated here (transcriptomics and proteomics/phosphoproteomics); other modalities such as metabolomics, genomics, and epigenomics can conceptually be integrated into the workflow. As more analytical frameworks for integrating multimodal data are created, we anticipate nanoSPLITS (and/or nanoSPLITS-derived technologies) will also enable greater insight into how different modalities interact with each other to orchestrate cell phenotypes in health and disease.

## Methods

### Reagents and chemicals

Deionized water (18.2 MΩ) was purified using a Barnstead Nanopure Infinity system (Los Angeles, CA, USA). n-dodecyl-β-D-maltoside (DDM), CDK1 inhibitor RO-3306, iodoacetamide (IAA), ammonium bicarbonate (ABC), and formic acid (FA) were obtained from Sigma (St. Louis, MO, USA). Nuclease-free water (not DEPC-treated), Trypsin (Promega, Madison, WI, USA) and Lys-C (Wako, Japan) were dissolved in 50 mM ABC before usage. Dithiothreitol (DTT, No-Weigh format), acetonitrile (ACN) with 0.1% FA, and water with 0.1% FA (MS grade) were purchased from Thermo Fisher Scientific (Waltham, MA, USA). SMART-Seq V4 Plus kit (Cat# R400753) was purchased from Takara Bio USA.

### Design, fabrication, and assembly of the nanoSPLITS chips

The nanoSPLITS chips were fabricated using standard photolithography, wet etching, and silanization as described previously^22^. Two different chips were designed and used in this study. Both contained 48 (4 x12) nanowells with a well diameter of 1.2 mm. The inter-well distance for the first chip was 2.5 mm while the second was 4.5 mm. Chip fabrication utilized a 25 mm x 75 mm glass slide pre-coated with chromium and photoresist (Telic Company, Valencia, USA). After photoresist exposure, development, and chromium etching (Transene), select areas of the chip were protected using Kapton tape before etching to a depth of ∼5 µm with buffered hydrofluoric acid. The freshly etched slide was dried by heating it at 120 °C for 1 h and then treated with oxygen plasma for 3 min (AP-300, Nordson March, Concord, USA). 2% (v/v) heptadecafluoro-1,1,2,2-tetrahydrodecyl-dimethylchlorosilane (PFDS, Gelest, Germany) in 2,2,4-trimethylpentane was applied onto the chip surface and incubated for 30 min to allow for silanization. The remaining chromium covering the wells was removed with etchant, leaving elevated hydrophilic nanowells surrounded by a hydrophobic background. To prevent retention of mRNA via interaction with free silanols on the hydrophilic surface of the nanowells, freshly etched chips were exposed to chlorotrimethylsilane under vacuum overnight to passivate the glass surface. A glass frame was epoxied to a standard glass cover slide so that it could be easily removed from the 2.5 mm inter-well distance chips for droplet splitting. For the 4.5 mm inter-well distance chips, PEEK chip covers were machined to fit the chip. Chips were wrapped in parafilm and aluminum foil for long-term storage and intermediate steps during sample preparation.

### Cell culture

Two murine cell lines (NAL1A clone C1C10 is referred to as C10 and is a non-transformed alveolar type II epithelial cell line derived from normal BALB/c mouse lungs; SVEC4-10, an endothelial cell line derived from axillary lymph node vessels) were cultured at 37°C and 5% CO_2_ in Dulbecco’s Modified Eagle’s Medium supplemented with 10% fetal bovine serum and 1× penicillin-streptomycin (Sigma, St. Louis, MO, USA). For the RO-3306 treatment, C10 cells were seeded at 200,000 cells per dish and incubated overnight. Treated cells were cultured with 10 µM RO-3306 for 36 h before harvesting. Control C10 cells were cultured similarly with vehicle (DMSO). The cultured cell lines were collected in a 15 ml tube and centrifuged at 1,000 × g for 3 min to remove the medium. Cell pellets were washed three times by PBS, then counted to obtain cell concentration. PBS was then added to achieve a concentration of ∼200 x 10^6^ cells/mL. Immediately before cell sorting, the cell-containing PBS solution was passed through a 40 µm cell strainer (Falcon™ Round-Bottom Polystyrene Test Tubes with Cell Strainer Snap Cap, FisherScientific) to remove aggregated cells.

Human islets (1,000-3000 islet equivalents) were obtained from Prodo Laboratories. Islets were immediately culture in PIM(S) media in a humidified incubator at 37°C and 5% CO2 after being received. After 24 hours in PIM(S) media, islets cells were collected and centrifuged at 100 x g for 5 mins and resuspended in TrypLE^TM^ for cell dissociation by incubating in a 37°C water bath for 15 to 30 minutes along with gentle pipetting until islets were visibly dispersed and no clumps of cells remained. The dissociated islet cells were centrifuge at 300 x g for 5 minutes; the supernatant was removed, and cells were resuspended in PBS. The dissociated cells were stained with 0.1 µM Calcein AM and 15 µM propidium iodide for cell viability prior to cell sorting with the cellenONE.

### CellenONE cell sorting and nanoSPLITS workflow

Before cell sorting, all nanoSPLITS chips were prepared by the addition of 200-nL hypotonic solution consisting of 0.1% DDM in 10 mM Tris to each nanowell. A cellenONE instrument equipped with a glass piezo capillary (P-20-CM) for dispensing and aspiration was utilized for single-cell isolation. Sorting parameters included a pulse length of 50 µs, a nozzle voltage of 80 V, a frequency of 500 Hz, a LED delay of 200 µs, and a LED pulse of 3 µs. The slide stage was operated at dew-point control mode to reduce droplet evaporation. Cells were isolated based on their size, circularity, and elongation in order to exclude apoptotic cells, doublets, or cell debris. For C10 cells, this corresponded to 25 to 40 µm in diameter, maximum circularity of 1.15, and maximum elongation of 2, while SVEC cells were 24 to 32 µm in diameter, maximum circularity of 1.15, and maximum elongation of 2. All cells were sorted based on bright field images in real time. The pooled C10 experiment had 11, 3, and 1 C10 cells sorted into each nanowell on a single 2.5 mm inter-well distance chip. For the SVEC and C10 comparison experiment, a single 48 well chip with 4.5 mm inter-well distance was used for each cell type and had a single cell sorted into each well. To perform the transferring identifications based on FAIMS filtering (TIFF) methodology for scProteomics^23^, a library chip was also prepared containing 20 cells per nanowell, with each cell type sorted separately on the same chip to reduce technical variation. After sorting, all chips were wrapped in parafilm and aluminum foil before being snap-frozen and stored at -80°C, which partially served to induce cell lysis via freeze-thaw. All associated settings, single-cell images, and metadata can be accessed at the GitHub repository provided (https://github.com/Cajun-data/nanoSPLITS).

To accomplish splitting of the cell lysate, chips were first allowed to thaw briefly on ice. For each split, a complementary chip was prepared that contained the same 200 nL of 0.1% DDM in 10 mM Tris on each nanowell. The bottom chip containing the cell lysate was placed on an aluminum chip holder that was pre-cooled to 4°C within a PCR workstation (AirClean Systems AC600). Precut 1/32” thick polyurethane foam was placed around wells on the exterior of this bottom chip while the top chip was slowly lowered onto the polyurethane foam (**<u>Movie S1</u>**). Wells were manually aligned for each chip before manual pressure was applied equally across the chip to merge the droplets for each chip. Pressure was held for 15 seconds before releasing. The droplets were merged twice more following this process. For consistency, the top chip was used for scRNAseq in all experiments while the bottom chip that initially contained the cell lysate was utilized in scProteomics (apart from the data generated for **<u>Fig. S3</u>**). After merging, the top chip was immediately transferred into a 96-well or 384-well UV-treated plate containing RT-PCR reagents. For the pooled C10 (11, 3, and 1 cell) experiment, the transfer was performed by adding 1µL of RT-PCR buffer to each nanowell before withdrawing the entire volume and adding it to a 96-well plate. For all other nanoSPLITS experiments, the transfer was accomplished by laying the 4.5 mm inter-well distance chip onto a 384-well plate containing wells with the RT-PCR mix, sealed with a PCR plate seal, and then centrifuged at 3,500 x g for 1 minute.

### Sample preparation and LC-MS/MS analysis for scProteomics

All post-split chips were first allowed to dry out before placing them into the humidified nanoPOTS platform for sample processing. Protein extraction was accomplished by dispensing 150 nL of extraction buffer containing 50 mM ABC, 0.1% DDM, 0.3 x diluted PBS, and 2 mM DTT and incubating the chip at 50°C for 90 min. Denatured and reduced proteins were alkylated through the addition of 50 nL 15 mM IAA before incubation for 30 min in darkness at room temperature. Alkylated proteins were then digested by adding 50 nL 50 mM ABC with 0.1 ng/nL of Lys-C and 0.4 ng/nL of trypsin and incubating at 37°C overnight. The digestion reaction was then quenched by adding 50 nL of 5% formic acid before drying the chip under vacuum at room temperature. All chips were stored in a -20°C until LC-MS analysis.

We employed the in-house assembled nanoPOTS autosampler for LC-MS analysis. The autosampler contains a custom packed SPE column (100 μm i.d., 4 cm, 5 μm particle size, 300 Å pore size C18 material, Phenomenex) and analytical LC column (50 μm i.d., 25 cm long, 1.7 μm particle size, 190 Å pore size C18 material, Waters) with a self-pack picofrit (cat. no. PF360-50-10-N-5, New Objective, Littleton, MA). The analytical column was heated to 50 °C using AgileSleeve column heater (Analytical Sales and services, Inc., Flanders, NJ). Briefly, samples were dissolved with Buffer A (0.1% formic acid in water) on the chip, then trapped on the SPE column for 5 min. After washing the peptides, samples were eluted at 100 nL/min and separated using a 60 min gradient from 8% to 35% Buffer B (0.1% formic acid in acetonitrile).

An Orbitrap Eclipse Tribrid MS (ThermoFisher Scientific) with FAIMS, operated in data-dependent acquisition mode, was used for all analyses. Source settings included a spray voltage of 2,400 V, ion transfer tube temperature of 200°C, and carrier gas flow of 4.6 L/min. For TIFF method^23^ samples, ionized peptides were fractionated by the FAIMS interface using internal CV stepping (-45, -60, and -75 V) with a total cycle time of 0.8 s per CV. Fractionated ions within a mass range 350-1600 m/z were acquired at 120,000 resolution with a max injection time of 500 ms, AGC target of 1E6, RF lens of 30%. Tandem mass spectra were collected in the ion trap with an AGC target of 20,000, a “rapid” ion trap scan rate, an isolation window of 1.4 *m/z*, a maximum injection time of 120 ms, and a HCD collision energy of 30%.

For the TIFF library samples, a single CV was used for each LC-MS run with slight modifications to the above method where cycle time was increased to 2 s and maximum injection time was set to 118 ms. Precursor ions with a minimum intensity of 1E4 were selected for fragmentation by 30% HCD and scanned in an ion trap with an AGC of 2E4 and an IT of 150 ms. Precursor ions with intensities > 1E4 were fragmented by 30% HCD and scanned with an AGC of 2E4 and an IT of 254 ms.

### RT-PCR, sequencing, and read mapping for scRNAseq

Following the transfer of samples into a 384-well plate containing RT-PCR buffer with 3’ SMART-Seq CDS Primer IIA (SMART-Seq® v4 PLUS Kit, cat# R400753), the samples were immediately denatured at 72°C for 3 min and chilled on ice for at least 2 min. Full length cDNA was generated by adding RT mix to each tube and incubating at 42°C for 90 min; followed by heat inactivation at 70°C for 10 min. 18 cycles of cDNA amplification were done to generate enough cDNA for template library according to SMART-Seq® v4 PLUS Kit instruction. The SMART-Seq Library Prep Kit and Unique Dual Index Kit (cat# R400745) were used to generate barcoded template library for sequencing. Single-read sequencing of the cDNA libraries with a read length of 150 was performed on NextSeq 550 Sequencing System using NextSeq 500/550 High Output v2 kit (150 cycles, cat#20024907). Data quality was assessed with FastQC and read-trimming was conducted using BBDuk. Reads were aligned to the mouse genome (Genome Reference Consortium Mouse Build 39) or human genome (Human genome assembly GRCh38) using STAR (https://github.com/alexdobin/STAR). BAM file outputs were mapped to genes using ‘featureCounts’ as part of the Subread package with default settings or using the -O, -M and --fraction settings to allow genes with overlapping features to be counted.

### Database searching and data analysis

All proteomic data raw files were processed by FragPipe^54^ (version 17.1 for C10 and SVEC cells, version 18.0 for G2/M arrested C10 cells, and version 20.0 for human islet cells)and searched against the *Mus musculus* or *Homo sapiens* UniProt protein sequence database with decoy sequences ( UP000000589 containing 17,201 forward entries, accessed 12/21 and UP000005640 containing 20,596 forward entries, accessed 05/23). Search settings included a precursor mass tolerance of +/-20 ppm, fragment mass tolerance of +/-0.5 Da, deisotoping, strict trypsin as the enzyme (with allowance for N-terminal semi-tryptic peptides), carbamidomethylation as a fixed modification, and several variable modifications, including oxidation of methionine, N-terminal acetylation, and S/Y/T phosphorylation. Protein and peptide identifications were filtered to a false discovery rate of less than 1% within FragPipe. For the TIFF methodology, IonQuant MBR (MBR) and MaxLFQ were set to “TRUE” and library MS datasets were assigned as such during the data import step. An MBR FDR of 5% at ion level was used to reduce false matching. FragPipe result files were then imported into RStudio (Build 461) for downstream analysis in the R environment (version 4.1.3). With regards to quality control filtering of samples in scProteomics and scRNAseq, thresholds were set based on the degree of protein or gene missingness. For scRNAseq data, a minimum of 5 read counts were used to filter genes with low abundance for all experiments with C10 and SVEC cells. For cases where complete data was needed, imputation was either performed with k-nearest neighbors imputation (n = 10 neighbors) with <60% missing values or by imputing from a normal distribution with a downshifted mean. For the pancreatic islet experiment, batch correction was performed with the ComBat.NA function from the MSnSet.utils R package. GO analysis was performed with the gprofiler2 R package and web application. Associated code for the data analysis are included in R markdown or script files at the nanoSPLITS GitHub repository (https://github.com/Cajun-data/nanoSPLITS).

## DATA AVAILABILITY

The mass spectrometry raw data have been deposited to the ProteomeXchange Consortium via the MassIVE partner repository with dataset identifiers MSV000089280, MSV000090828, and MSV000093330 (FTP Password: Nano4108, Nano4468, and Nano5238, respectively). The raw RNAseq data has been deposited to the Gene Expression Omnibus (GEO) under the identifiers GSE201575, GSE219047, and GSE247519.

## CODE AVAILABILITY

All scripts, result tables, single-cell images, and metadata used for figure generation are available at the nanoSPLITS GitHub repository (https://github.com/Cajun-data/nanoSPLITS).

## ACKNOWLEDGMENTS

We thank Drs. Clayton Mathews and Jing Chen at the University of Florida for providing a detailed protocol for culturing human islets, and Dr. Matthew Monroe for his assistance in depositing the raw proteomic data into MassIVE, as well as Nathan Johnson for the illustration utilized in Figure 1. This work was supported by a Laboratory Directed Research and Development award (I3T) from Pacific Northwest National Laboratory (Y.Z.). A portion of this research was performed on a National Institutes of Health (NIH) Common Fund, Human Biomolecular Atlas Program (HuBMAP) grant UH3CA256959 (L.P-T.). A portion of this research was also performed on a project award (10.46936/prtn.proj.2021.51834/60000340) from the Environmental Molecular Sciences Laboratory (J.C.B. and L.P-T.), a DOE Office of Science User Facility sponsored by the Biological and Environmental Research program under Contract No. DE-AC05-76RL01830.

## AUTHOR CONTRIBUTIONS

Y.Z., J.M.F., L.P-T., and L.M.M. conceptualized and designed the research. J.M.F. and S.M.W. fabricated nanoSPLITS microchips. K.M.E., L.M.M., J.M.F., J.C.B., W.J.Q. and Y.Z. performed the cell culture, cell sorting, and proteomic sample preparation. J.M.F, D.J.D., and L.M.B. performed data analysis. S.M.W., J.M.F., and R.J.M. performed the LC-MS analysis. L.M.M. and H.D.M. performed the RNAseq and associated read mapping. J.M.F., Y.Z., and L.M.M. wrote the manuscript. J.M.F., Y.Z., L.P-T., J.C.B., A.S., W.J.Q., J.W.B., and L.M.M. edited the manuscript. The final manuscript was read and approved by all authors.

## COMPETING INTERESTS

J.C.B., J.W.B, and A.S. are employees of Scienion/Cellenion. Y.Z. is an employee of Genentech Inc. and shareholder of Roche Group. Battelle Memorial Institute has submitted a U.S. patent application for the design of nanoSPLITS devices and the associated operation methods (Application number: 17/954,834, filed 09/28/2022; Inventors: Y.Z., J. M. F., L.M.M., and L.P.T.; Status of application: Pending). Other authors declare no other competing interests.

## Notes

### Summary of Updates

We added new experiments to apply the technology in primary human cells and developed an efficient approach for facile identification of unknown cell types, and detecting their protein markers by mapping transcriptomic data to existing large-scale single-cell RNA sequencing reference databases.

