## Supplementary information for "Parallel measurement of transcriptomes and proteomes from same single cells using nanodroplet splitting"

**Table of Contents:**


| Title | Page |
| --- | --- |
| Figure S1–Lysis buffer optimization results | 4 |
| Figure S2– Imaging of droplet array after nanoSPLITS | 5 |
| Table S1– Quantification of fluorescence intensities | 6 |
| Figure S3– Evaluation of post-split protein recovery with proteomics | 7 |
| Figure S4– Pearson correlations for 11 and 1 C10 cell | 8 |
| Figure S5– Distributions of correlations and overlapping identifications (SVEC and C10 cells) | 9 |
| Figure S6– Histogram of global across-modality correlations and the distribution of genes by TPM | 10 |
| Figure S7– PCAs of C10 and SVEC from scProteomics and scRNAseq | 11 |
| Figure S8– UMAPs of C10 cells with cell cycle marker genes and proteins | 12 |
| Figure S9– Cell size distributions for treated and control C10 cells | 13 |
| Figure S10–PCAs and WNN-UMAP of G2/M arrested and untreated C10 cells | 14 |
| Figure S11–scRNAseq clustergram of G2/M arrested and untreated C10 cells | 15 |
| Figure S12–Covarying proteins identified in cluster 3 related to cell cycle | 16 |
| Figure S13–Protein-gene pairs concordant between scRNAseq and scProteomics | 17 |
| Figure S14– UMAP feature maps of established islet protein markers | 18 |
| Figure S15– PCAs of islet cells after spls-DA classification and Seurat annotation of scProteomics data | 19 |
| Figure S16– Flowchart for annotation of human pancreatic islet cells | 20 |
| Figure S17– Clustergram of protein-protein correlations across alpha cells | 21 |
| Figure S18– Clustergram of protein-protein correlations across beta cells | 22 |
| Figure S19– Histogram of mRNA-protein correlations for genes/proteins quantified in both modalities | 23 |
| Figure S20–Gene ontological analyses of cluster 4 subcluster | 24 |
| Figure S21: Pearson correlations between PSAP/GRN and CTSA/CTSB | 25 |

Figure S22–Overlap of proteins/genes increasing or decreasing in abundance 26

Figure S23– GO dotplot of top 4 enriched terms for three annotation sets 27

Figure S24– GO dotplot of enriched terms related to cell cycle or mitosis 28

Figure S25- GO terms of differentially abundant proteins and genes 29

Figure S26- Correlations between selected proteins involved in mitosis 30

Supplementary Results and Discussion 31-33

Supplementary Methods 34-36

References 37

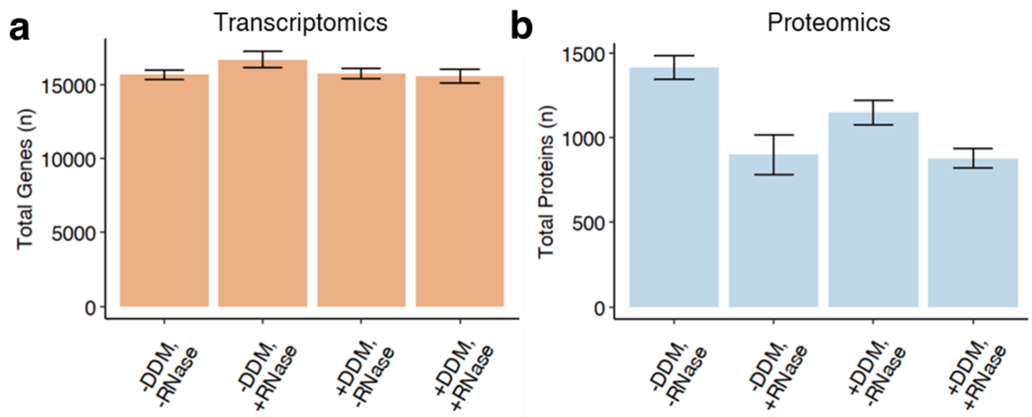


**Figure S1**: (**a)** Mean numbers of genes detected per condition with RNAseq (n = 2 for each condition) (**b**) Mean numbers of proteins detected per condition with label-free proteomics (n = 4 for each condition). Error bars represent +/- sd. Conditions in (**a**) and (**b**) indicated by “+” or “-" represent the presence or absence of 0.1 %DDM or 1 x RNase inhibitor, respectively.

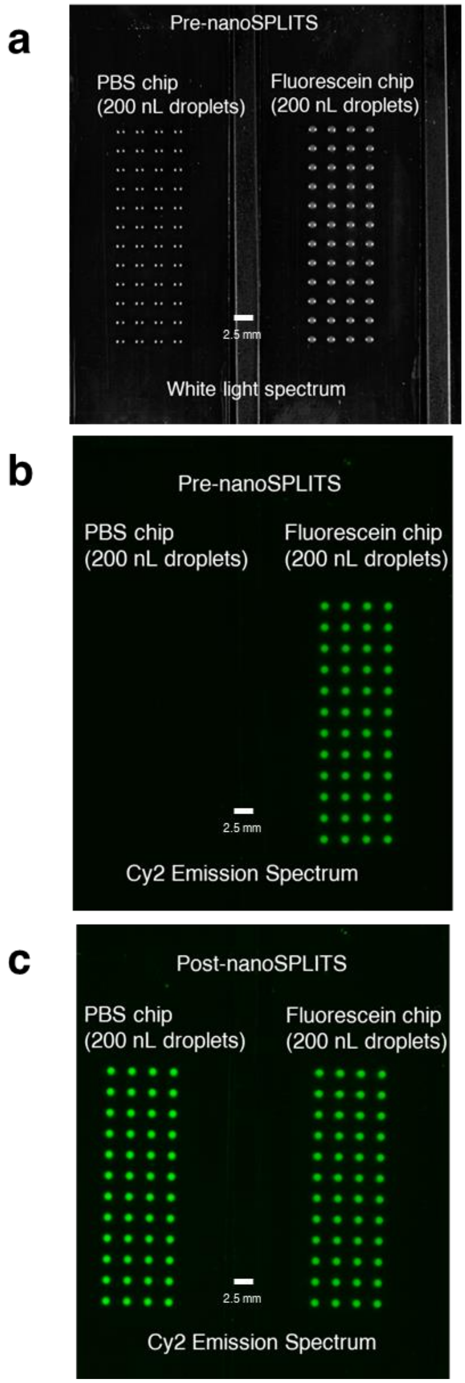


**Figure S2**: (**a**) Brightfield image of two nanoSPLITS chips before performing the merging and splitting process. The left chip contains 200 nL of phosphate-buffered saline (PBS) while the right chip contains 200 nL of 0.01% fluorescein in PBS (n = 48). (**b**) Same as (**a**) but filtered for Cy2 emission. (**c**) Cy2 emission image of the same two nanoSPLITS chips immediately after merging and splitting. White scale bar indicates a 2.5 mm distance, which is equivalent to the length between each nanoSPLIT droplet. Quantification of fluorescence was performed with ImageJ version 1.5.

| nanoSPLITS chip column # | Mean Column Fluorescence Intensity in PBS Chip (RFU) | Mean Column Fluorescence Intensity in fluorescein chip (RFU) | Split Ratio (Relative to fluorescein chip) |
|---|---|---|---|
| 1 | 4236.85 | 3421.87 | 45% |
| 2 | 3979.04 | 3499.85 | 47% |
| 3 | 4396.41 | 3584.41 | 45% |
| 4 | 4262.54 | 3511.98 | 45% |
| 5 | 4414.46 | 3483.15 | 45% |
| 6 | 4285.49 | 3464.50 | 45% |
| 7 | 4488.26 | 3503.46 | 44% |
| 8 | 4565.42 | 3713.30 | 45% |
| 9 | 4543.61 | 4149.19 | 48% |
| 10 | 4569.14 | 4178.50 | 48% |
| 11 | 4595.23 | 4141.59 | 47% |
| 12 | 4550.56 | 4197.75 | 48% |

**Table S1**: Quantification results after merging and splitting of chip containing 5,6-carboxyfluorescein and chip containing PBS. Post-split Cy2 emission images were imported into ImageJ for analysis. Each column represents an average of the four wells (n = 48 total).

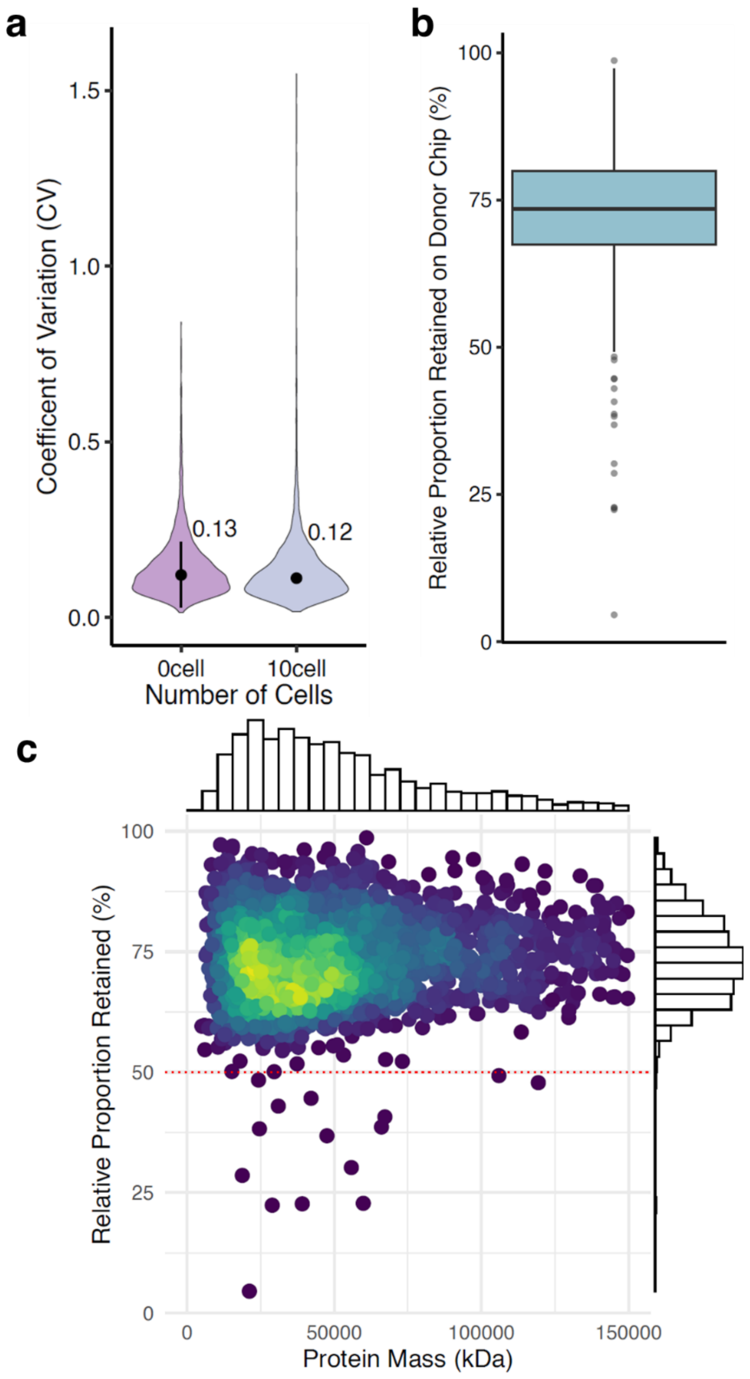


**Figure S3**: (**a**) Coefficients of variation (CV) of quantified protein intensities for droplet samples from the donor chip (10 C10 cells, n = 6) and the acceptor chip (0 C10 cells, n = 6) after splitting. (**b**) Boxplot showing the relative proportions of proteins retained on the donor chip (**c**) Scatter plot showing the relative proportion of protein retained on the donor chip (y-axis, defined as the mean protein intensity for each protein from the donor chip divided by the mean total intensity) and the protein molecular weight (x-axis). Density is indicated by the color and also displayed with marginal histograms along each axis.

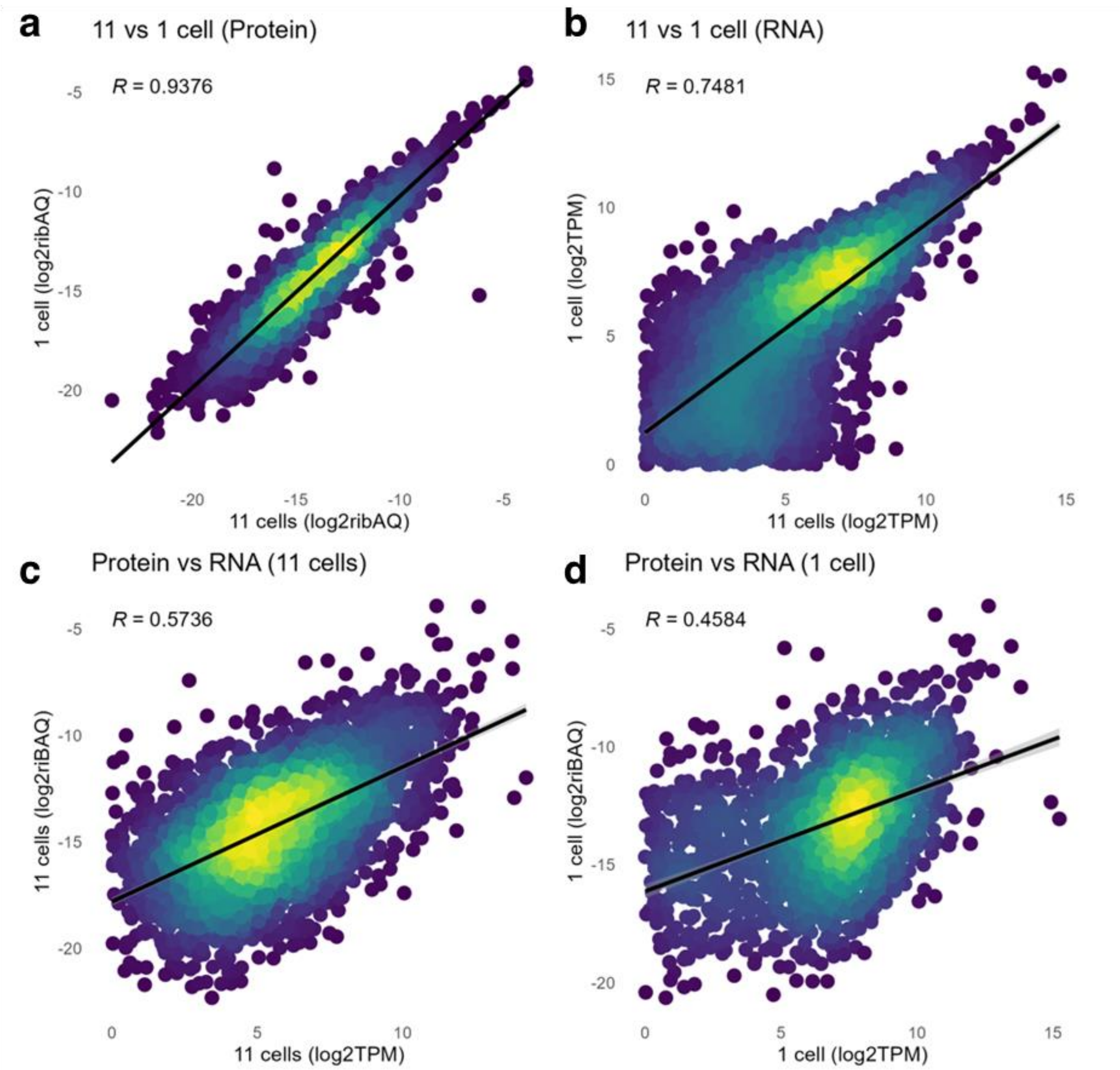


**Figure S4**: (**a**) Scatter plot of the correlations between the mean protein abundances for 11 C10 cells (n = 6, x-axis) and 1 C10 cell (n = 7, y-axis). Units are $log_2$-transformed relative intensity-based absolute quantification for proteomics (riBAQ) values. (**b**) Scatter plot of the correlation between the mean gene transcript abundances for 11 C10 cells (n = 6, x-axis) and 1 C10 cell (n = 7, y-axis). Units are $log_2$ of transcripts per million (TPM). (**c**) Scatter plot of the correlation between the mean protein abundance (y-axis) and mean gene transcript abundance (x-axis) for 11 C10 cells (n = 6). (**d**) Scatter plot of the correlation between the mean protein abundance (y-axis) and mean gene transcript abundance (x-axis) for 1 C10 cell (n = 7). Color indicates density of data points for all plots.

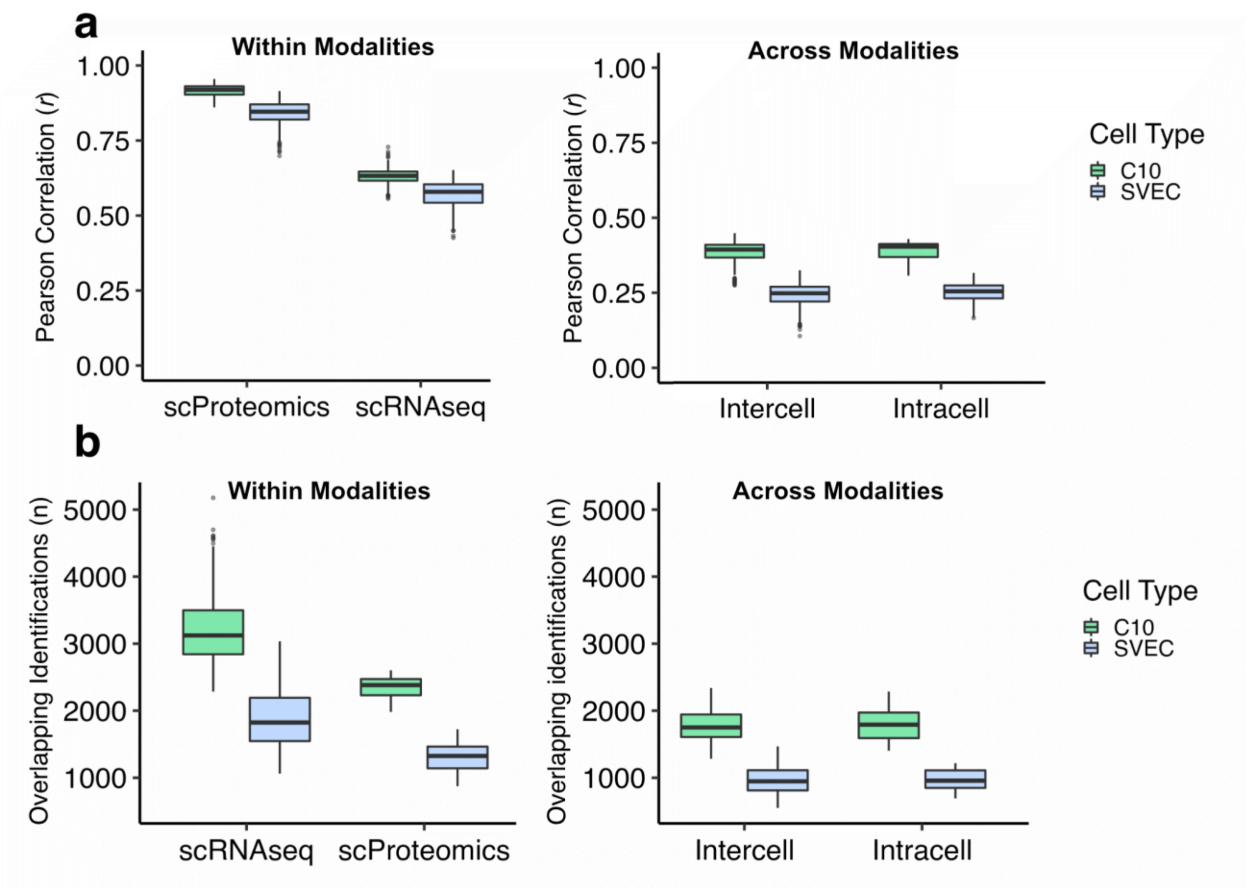


**Figure S5.** (**a**) Box plots showing the distributions of Pearson correlations, separated by cell types (C10 and SVEC) and modalities (scProteomics and scRNAseq). (**b**) Overlap in gene and protein identifications within and across modalities.

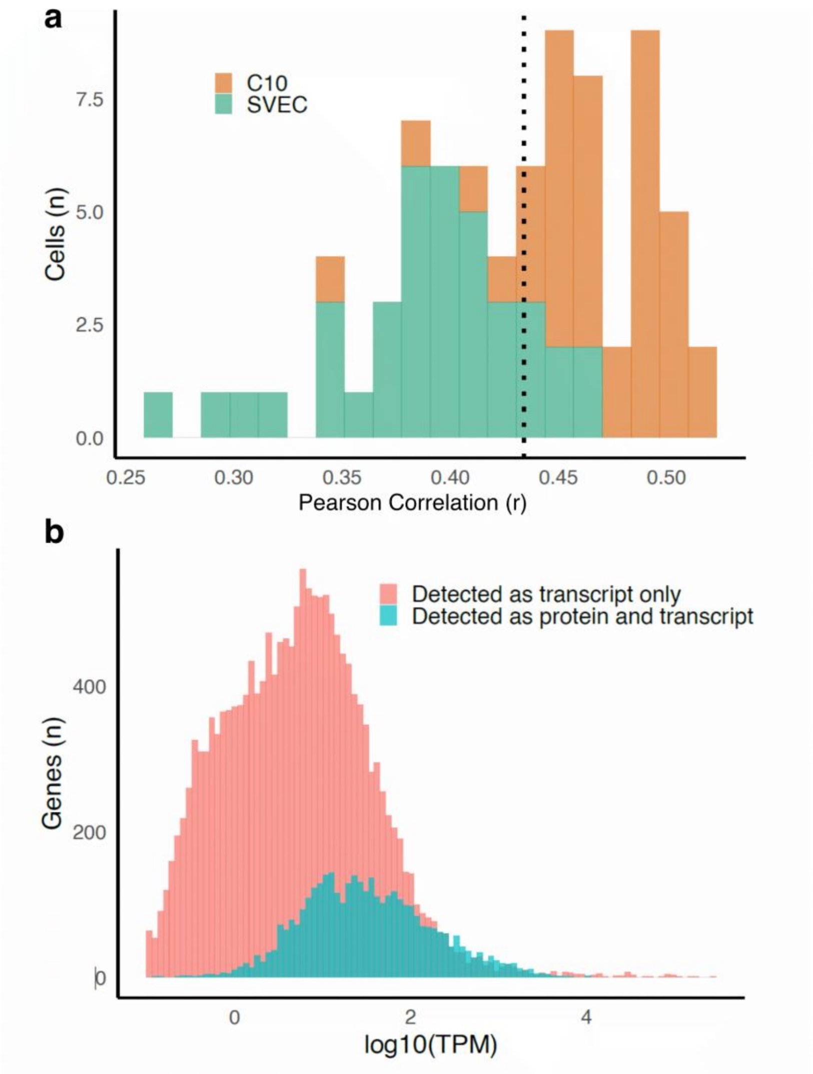


**Figure S6**: (**a**) Histogram of global across-modality correlations for C10 and SVEC cells. Color indicates cell type. The median (0.44) of the combined distribution is indicated with a dashed line. (**b**) Histogram of all genes detected in this study with transcripts per million (TPM) > 0.1 at mRNA and/or protein level, arranged by the average $\log_{10}$(TPM) of each gene detected.

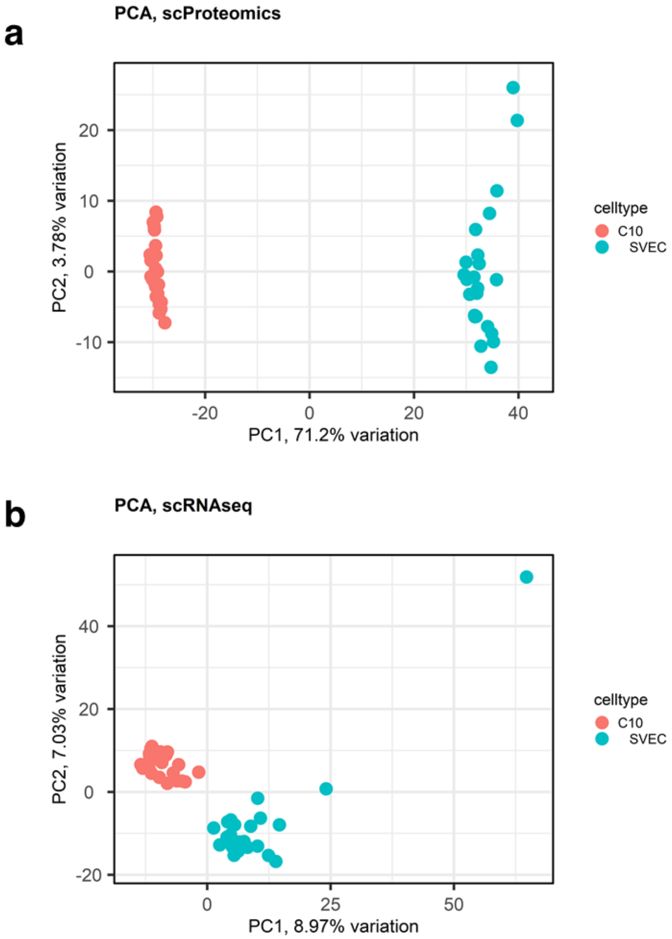


**Figure S7**: (**a**) PCA of SVEC (n = 23) and C10 (n = 26) cells using scProteomics data and (**b**) scRNAseq data.

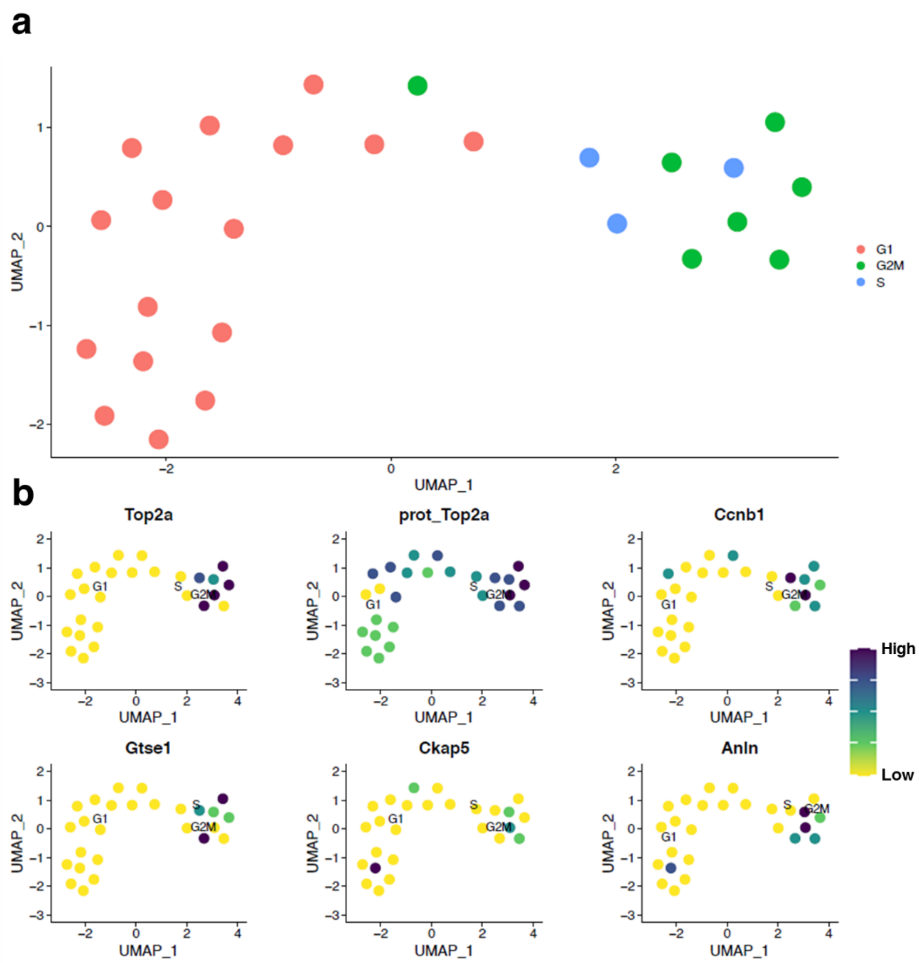


**Figure S8**: (**a**) UMAP generated by Seurat using known cell cycle features and colored by cell cycle (n = 26). (**b**) UMAP feature maps showing abundances of represented genes (and the protein TOP2A) grouped by assigned cell cycle phase. All expression values are Z-scores after scaling and centering of data.

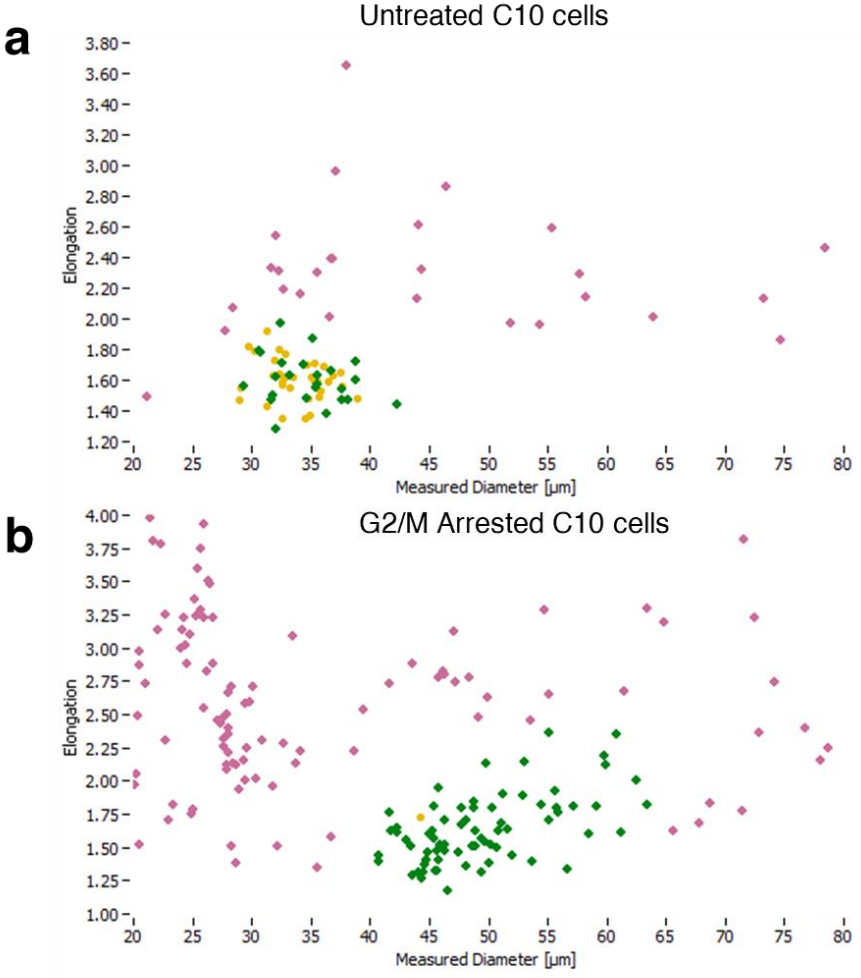


**Figure S9**: Scatter plot of untreated (control) (**a**) and RO-3306 treated (**b**) C10 cells sorted using the cellenONE. X-axis represents the measured diameter of the cell, while Y-axis represents the elongation of the cell (arbitrary value). Pink represents cells or debris that were excluded during sorting, yellow represents cells that were the correct size but not sorted due to more cells being found within the sedimentation/ejection zone, and green represents cells that were sorted.

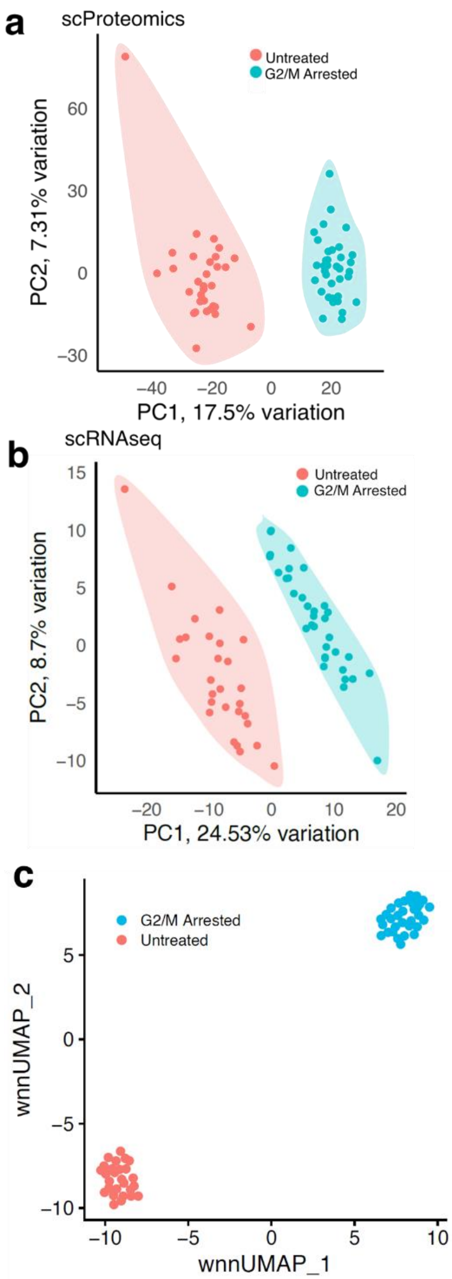


**Figure S10**: (**a**) scProteomics PCA (2,863 proteins with < 50% missing values) of untreated (n = 32) and G2/M arrested C10 cells (n = 36). (**b**) scRNAseq PCA (333 genes with complete data) of untreated control (n = 28) and G2/M arrested C10 cells (n = 34). (**c**) Integration of scProteomic and scRNAseq data with WNN UMAP generated using Seurat (1,722 mRNA-protein pairs, n = 31 for G2/M arrested and n = 27 for untreated).

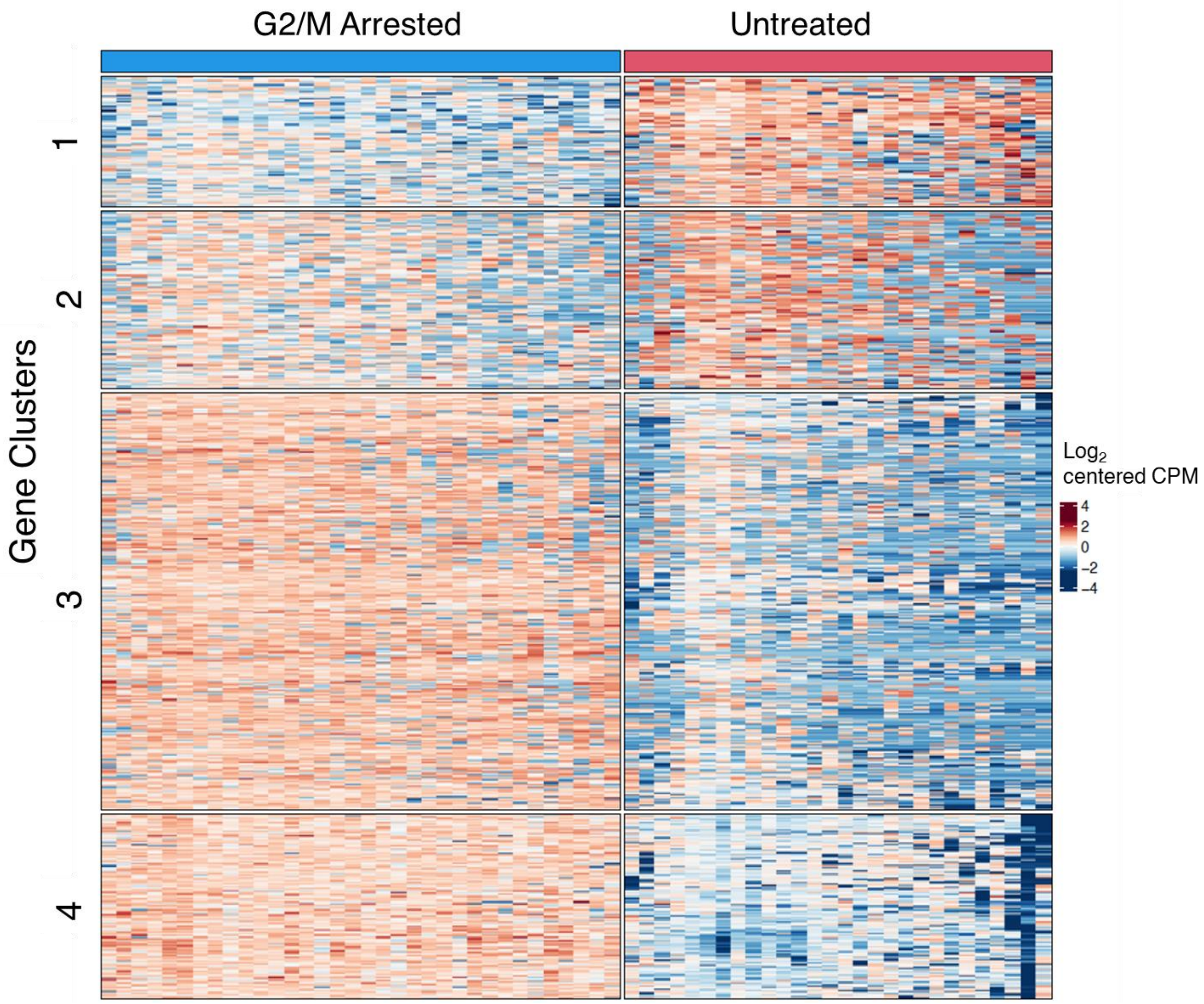


**Figure S11**: Clustergram of $\log_2$ centered counts per million (CPM) for differentially expressed genes between control (column cluster 1) and RO-3306 treated (column cluster 2) cells using scRNAseq data with adjusted *p*-values <= 0.01 and log2FC of > 1 or < -1 (830 genes).

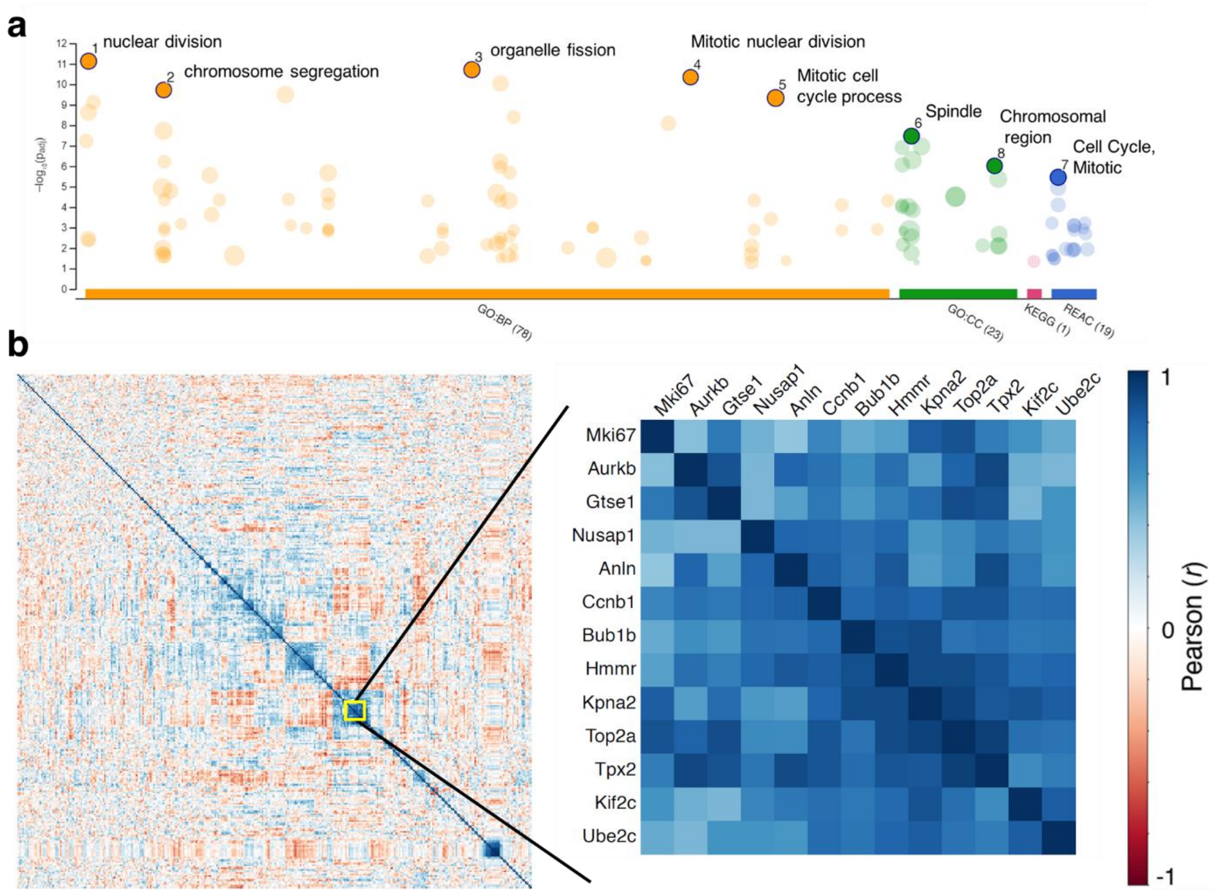


**Figure S12**: (**a**) Gene ontology enrichment of proteins found in cluster 3 of the differentially expressed scProteomic data (13 proteins total).(**b**) Left panel**;** Pearson correlation heatmap of all 327 differentially abundant proteins found in G2/M arrested cells. Right panel; enlarged area from the heatmap showing the 13 proteins from cluster 3.

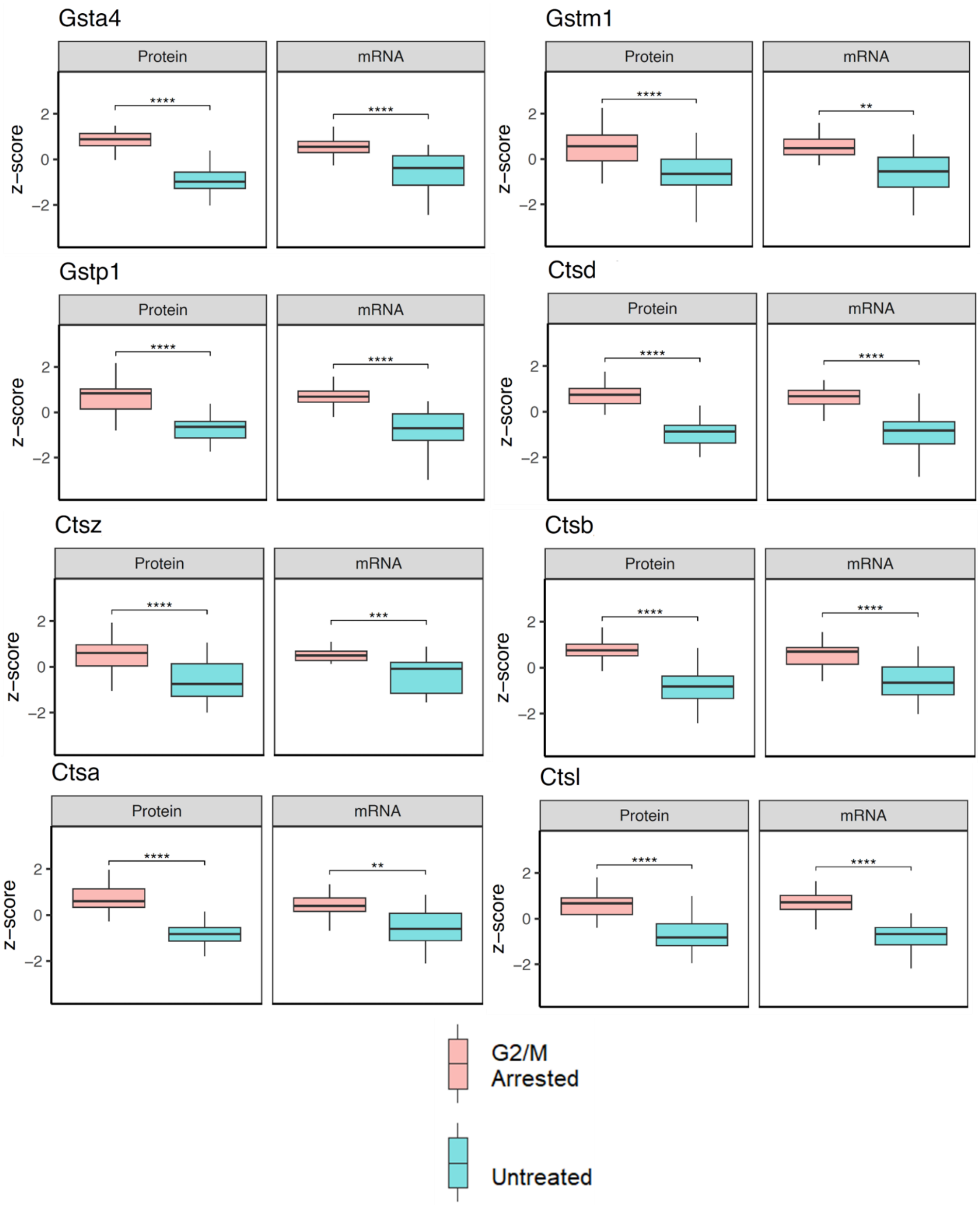


**Figure S13**: Boxplots of differentially abundant mRNA transcripts and proteins showing concordance between the two modalities, including several glutathione S-transferases (GSTA4, GSTM1, and GSTP1) and cathepsins (CTSA, CTSB, CTSD, CTSL, and CTSZ). Asterisks indicate the degree of significance after significance testing with Wilcoxon test (**** indicates FDR < 0.0001, *** FDR < 0.001 and ** FDR < 0.01).

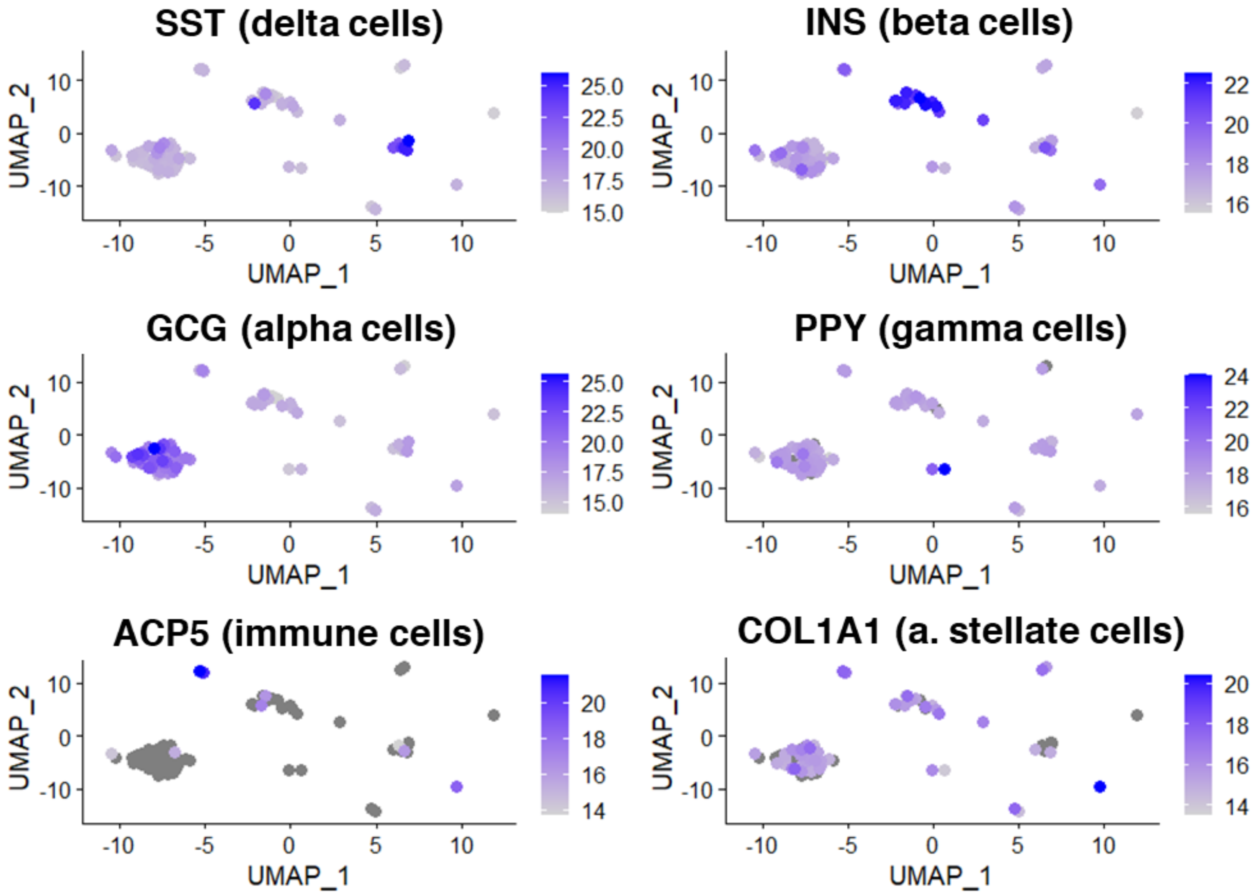


**Figure S14**: scProteomic UMAPs of established markers for several pancreatic islet cell types. Scale bars indicate log2(Intensity) for each protein separately.

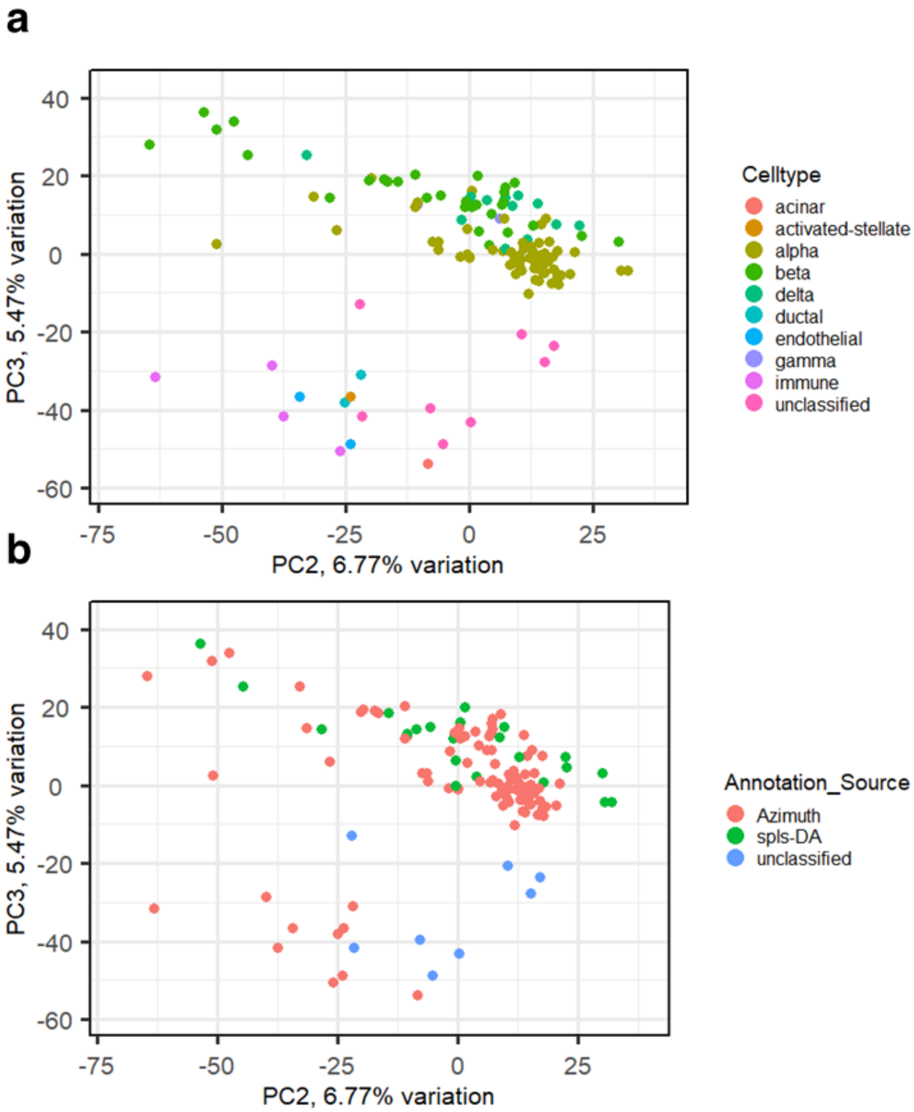


**Figure S15**: (**a**) PCA of the 126 pancreatic islet cells analyzed with scProteomics, colored by cell types (**b**) PCA of the same 126 cells but reflecting the source of our cell annotations (Azimuth, spls-DA, or "unclassified")

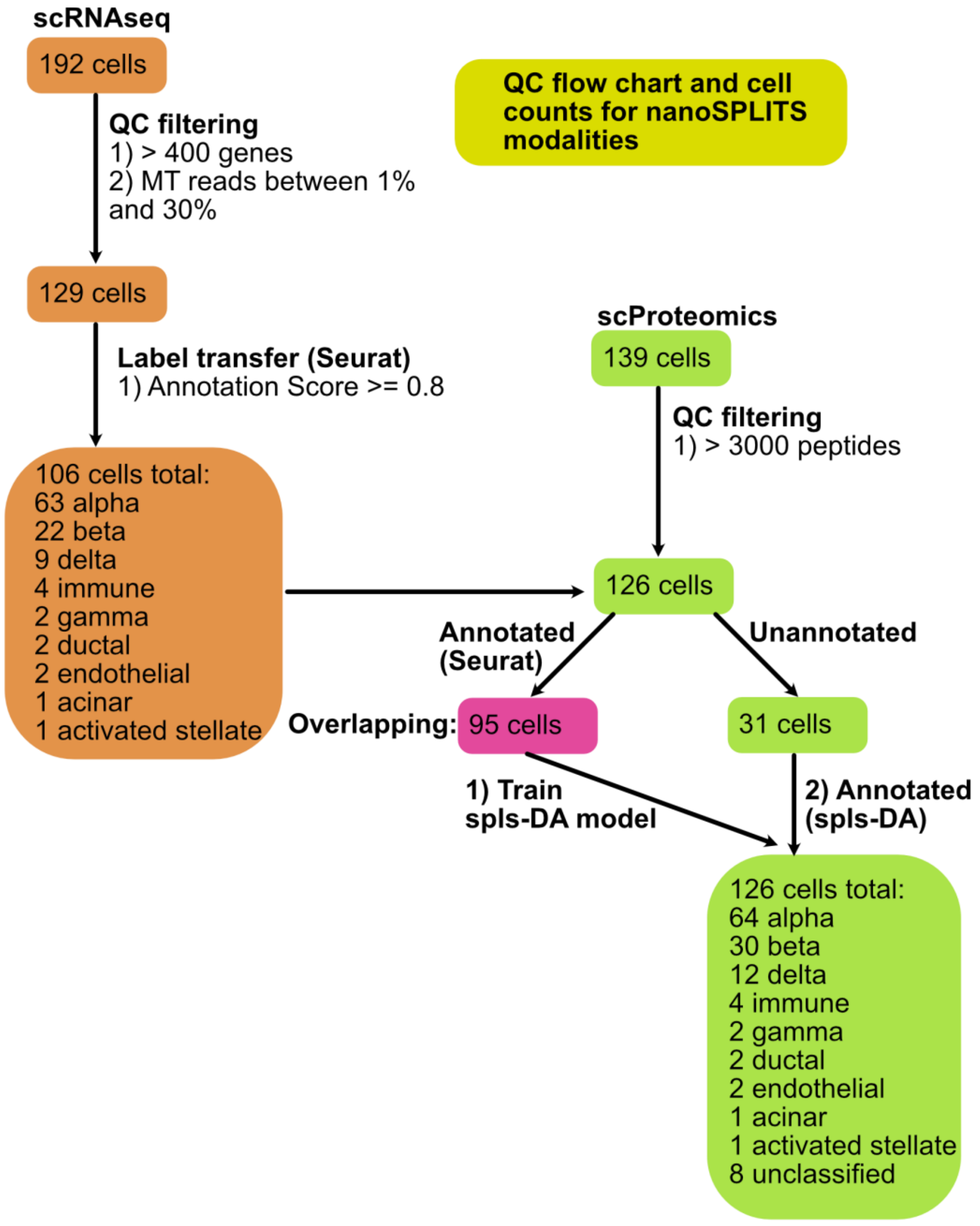


**Figure S16**: Flowchart for documenting the number of cells retained across quality control (QC) and annotation with Seurat and spls-DA.

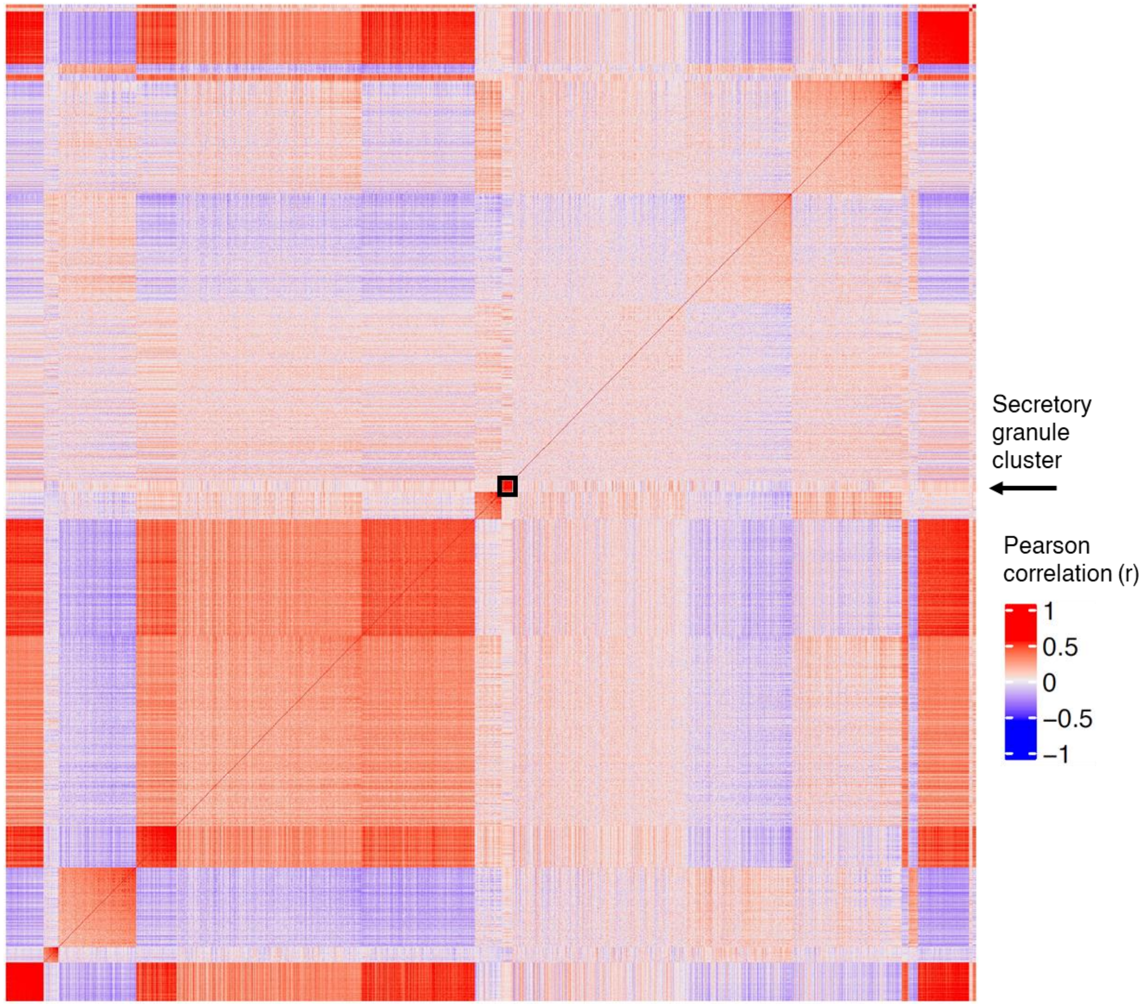


**Figure S17**: Clustergram of protein-protein Pearson correlations across all alpha cells. Rows and columns are individual proteins arranged using Gaussian mixture modeling (GMM) clusters and cluster member’s uncertainty values

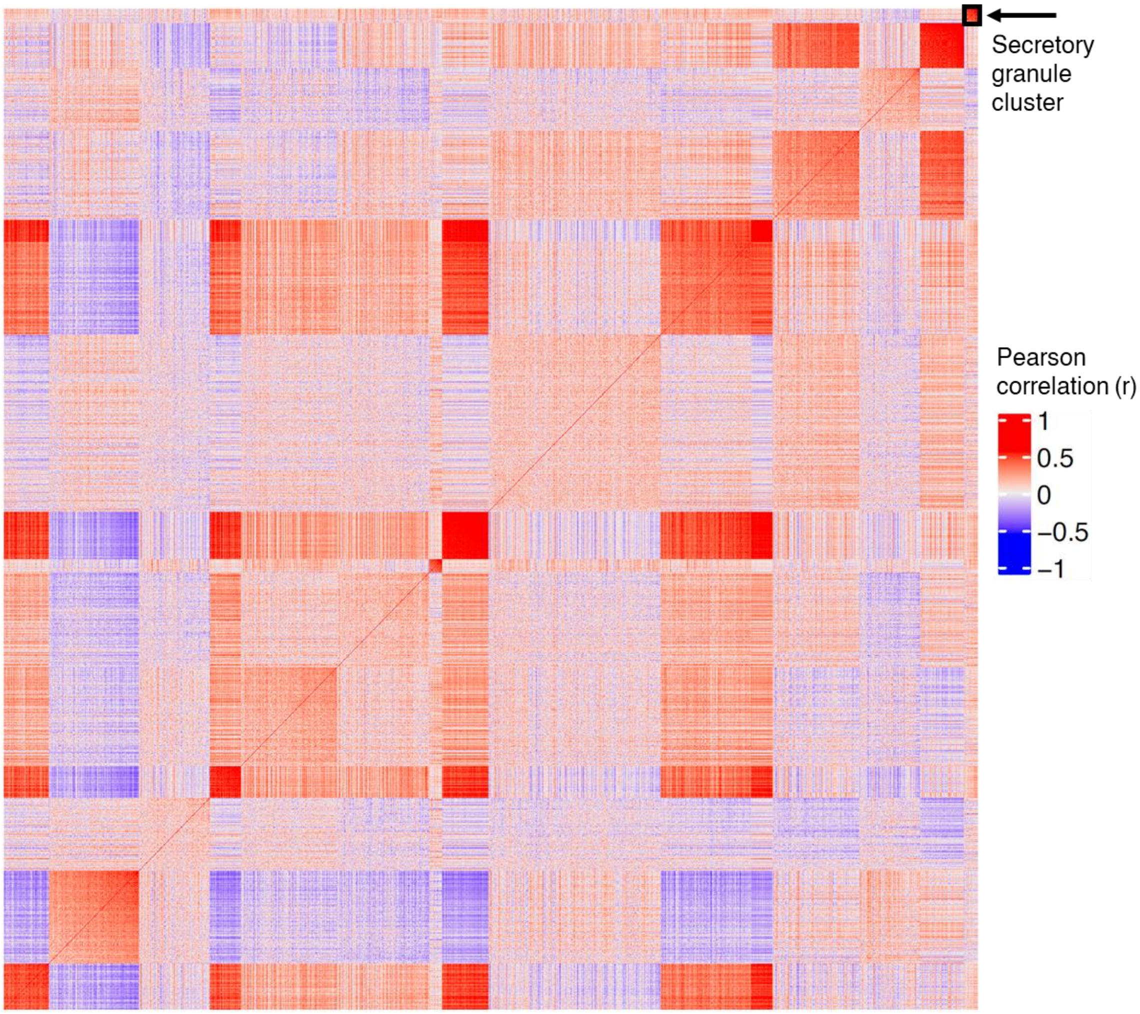


**Figure S18**: Clustergram of protein-protein Pearson correlations across all beta cells. Rows and columns are individual proteins arranged using GMM clusters and cluster member's uncertainty values.

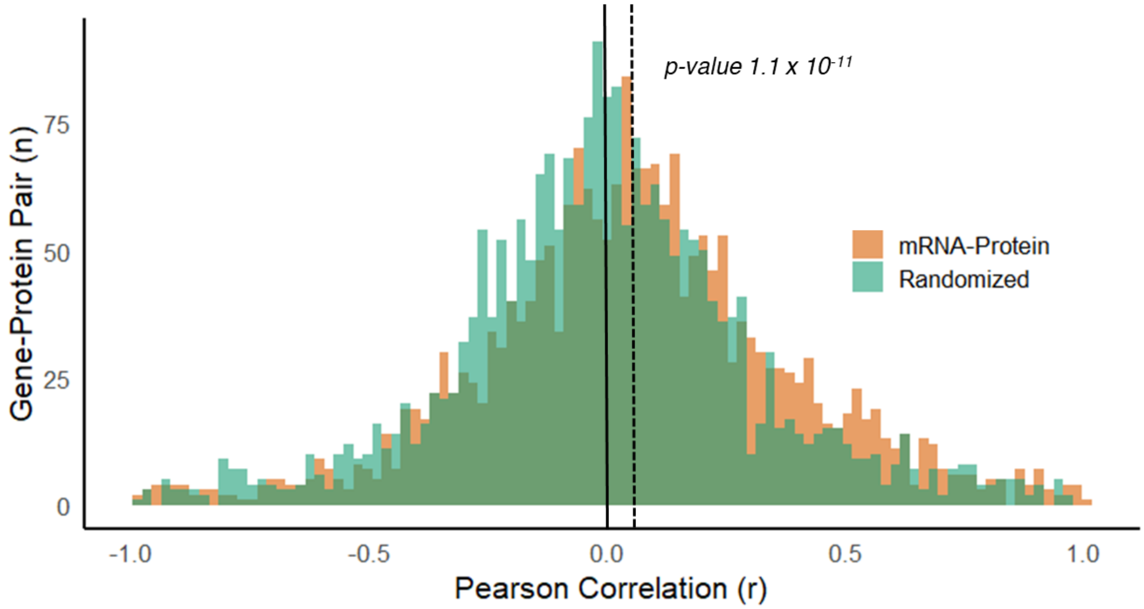


**Figure S19**: Histogram of mRNA-protein correlations for each gene quantified in both modalities with at least 4 observations for all pancreatic cell types. Statistical testing was performed by non-parametric Mann-Whitney. Solid and dashed lines indicate medians of randomized and real correlations, respectively.

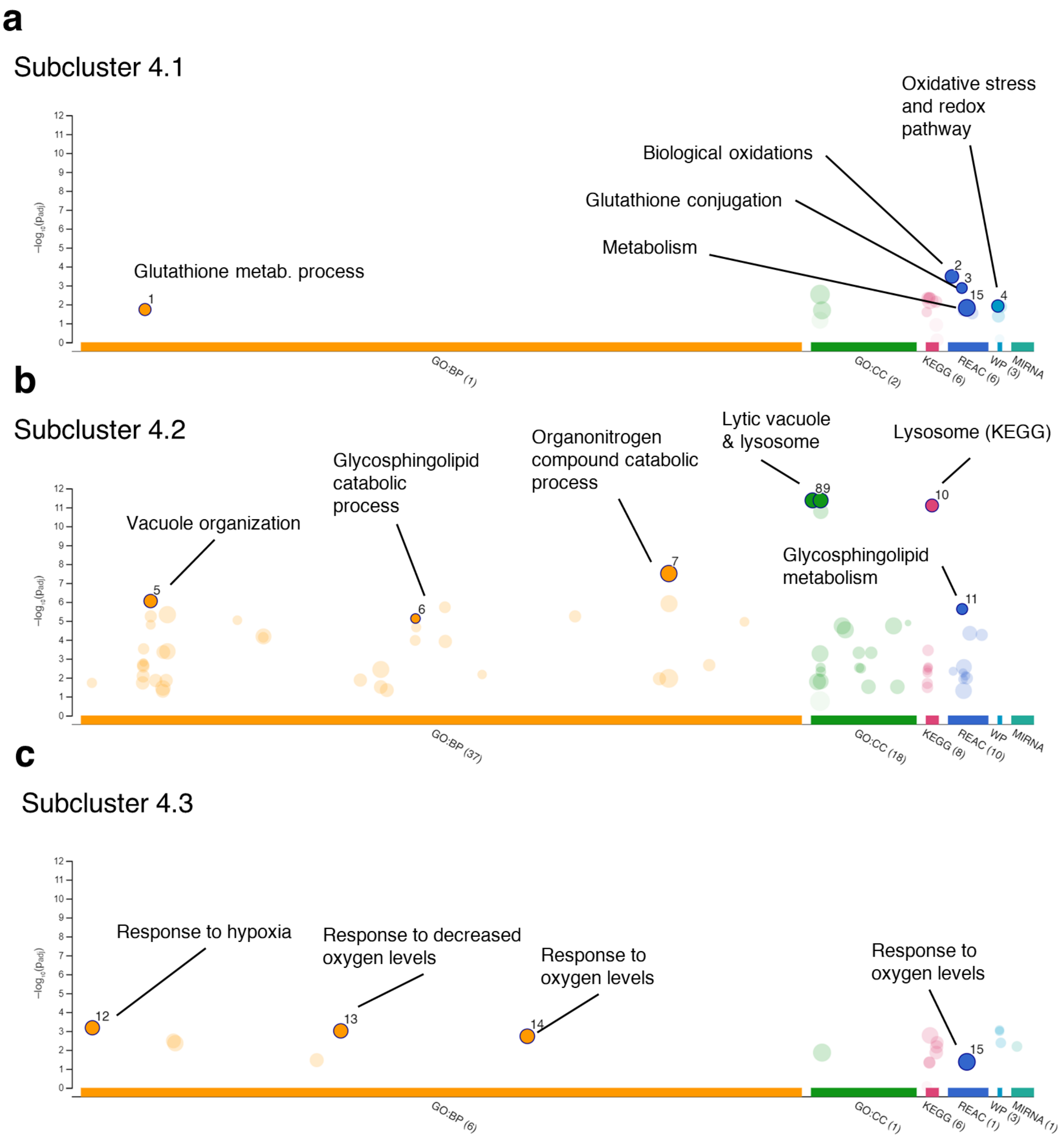


**Figure S20:** (**a**),(**b**), and (**c**) Gene ontology enrichment of subclusters 4.1 (17 proteins), 4.2 (8 proteins), and 4.3 (7 proteins), respectively. Selected terms are highlighted, with color indicating the domain from which the terms were assigned.

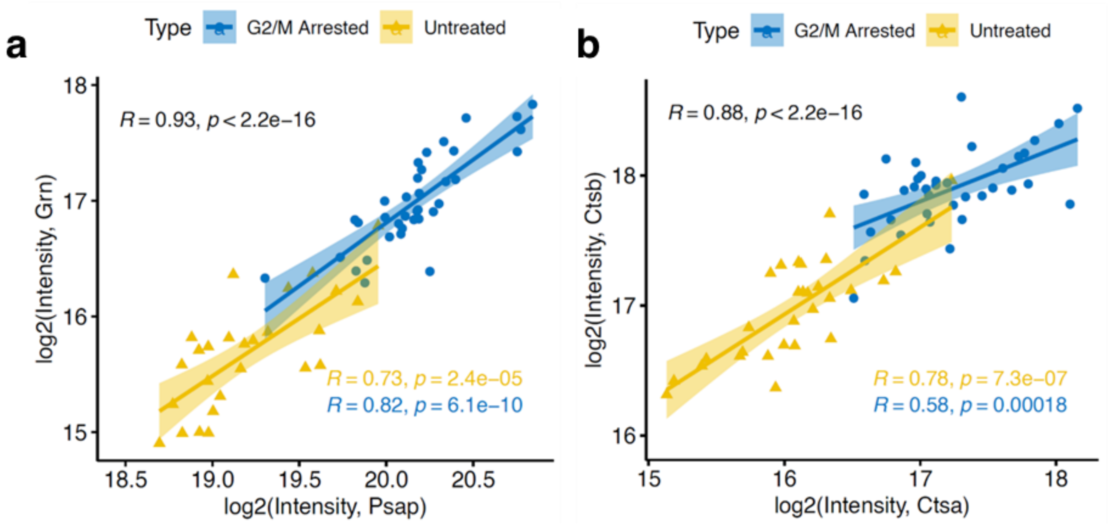


**Figure S21**: (**a**) Pearson correlations between PSAP and GRN using $\log_2$(Intensity). Confidence intervals and data points are colored based on the cell group (G2/M arrested or untreated). Color also indicates Pearson correlation and p-value for each group, with black (upper left) representing the combined correlation of all cells in the plot. (**b**) Same as (**a**) except comparing CTSA with CTSB.

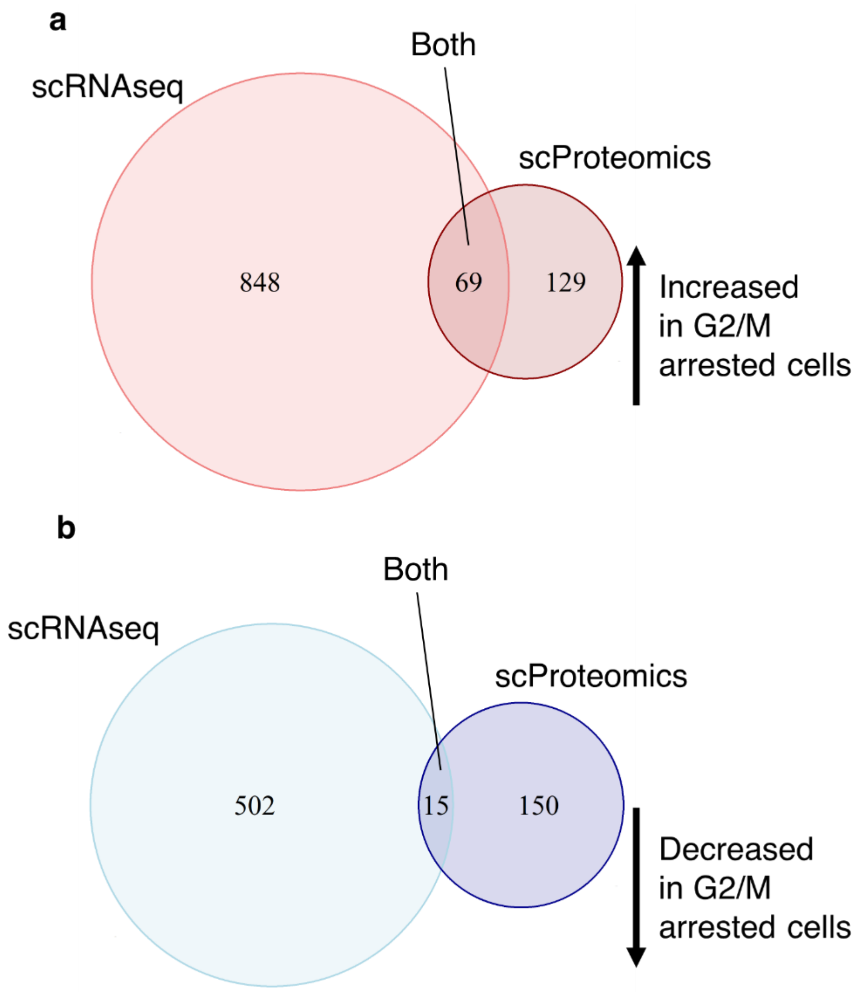


**Figure S22**: (**a**) Venn diagram of all genes and proteins significantly increased in abundance in G2/M arrested cells (FDR < 0.01 and $\log_2$FC > 0.5 or < -0.5) (**b**) Venn diagram of all genes and proteins significantly decreased in abundance in G2/M arrested cells (FDR < 0.01 and $\log_2$FC > 0.5 or < -0.5)

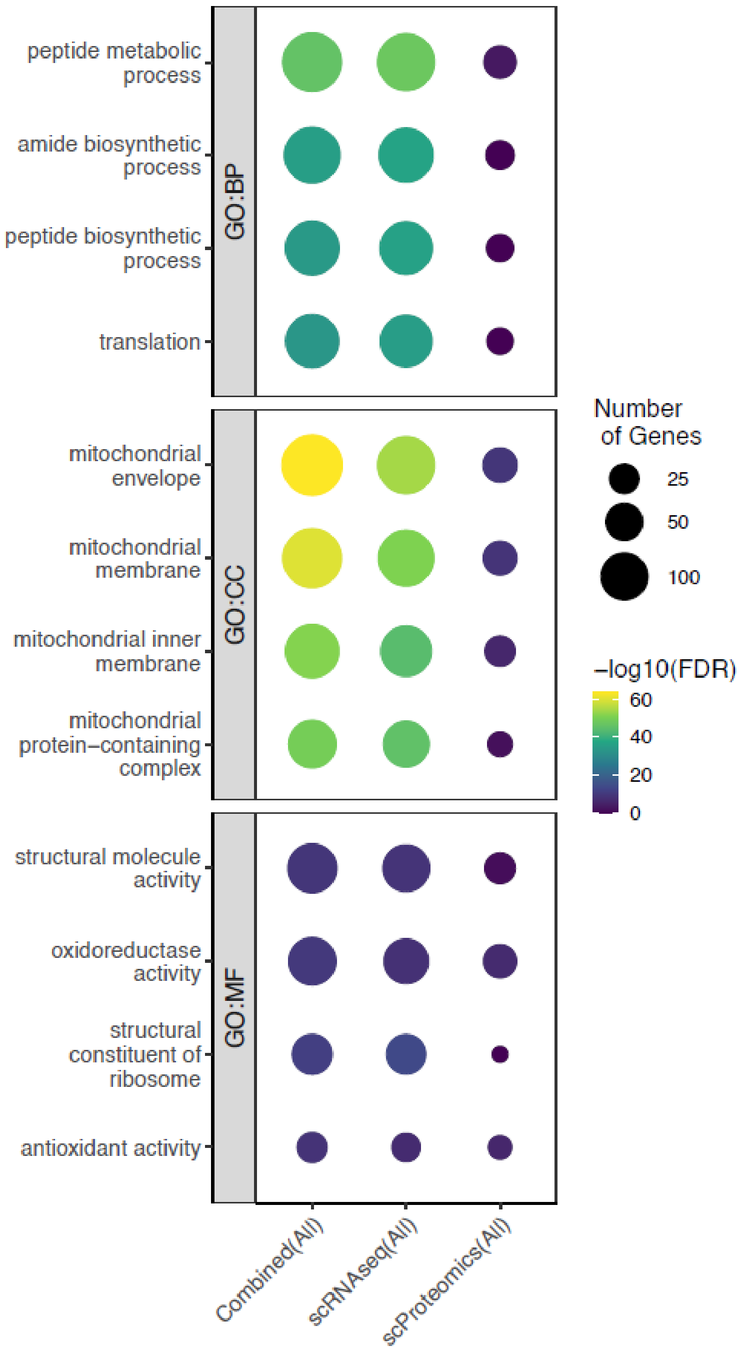


**Figure S23**: Dotplot of the top 4 most significant GO terms for each annotation set (GO:MF, GO:CC, and GO:BP). "All" represents all genes or proteins with $log_2FC$ +/- 0.5 and FDR < 0.01. "Combined" refers to the combination of significant genes and proteins from scRNAseq and scProteomics.

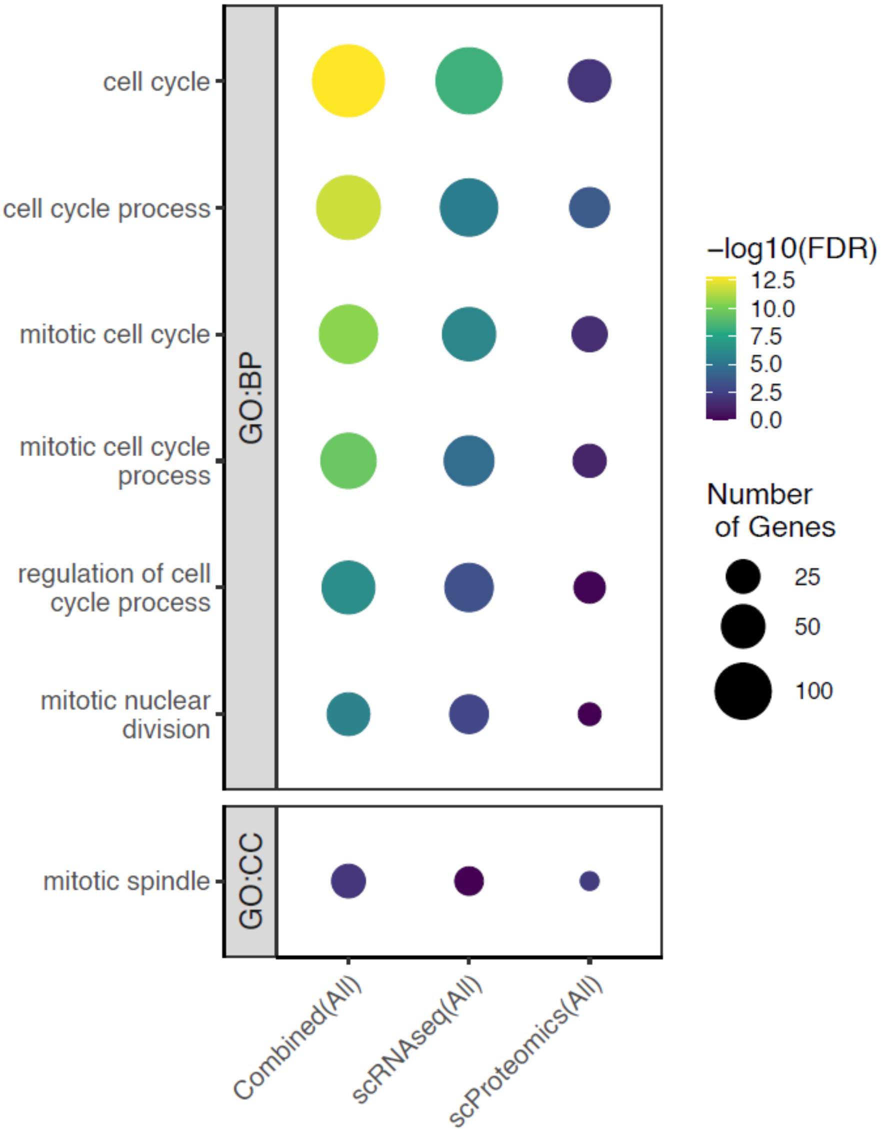


**Figure S24**: Dotplot of all terms related to cell cycle or mitosis found enriched. “All” represents all genes or proteins with $log_2FC$ +/- 0.5 and FDR < 0.01. “Combined” refers to the combination of significant genes and proteins from scRNAseq and scProteomics.

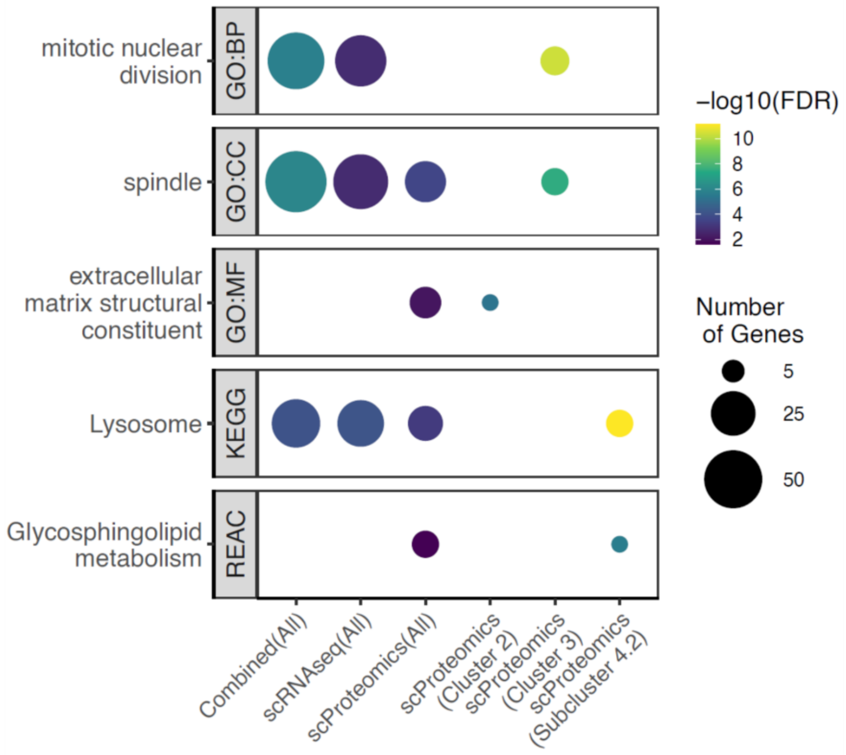


**Figure S25**: Enrichment of GO terms (y-axis) for differentially abundant proteins and genes found in the scProteomic and scRNAseq data (x-axis) with FDR < 0.01 and log2FC of +/- 0.5. Point size represents the number of genes (proteins) and color represents -log10(FDR)

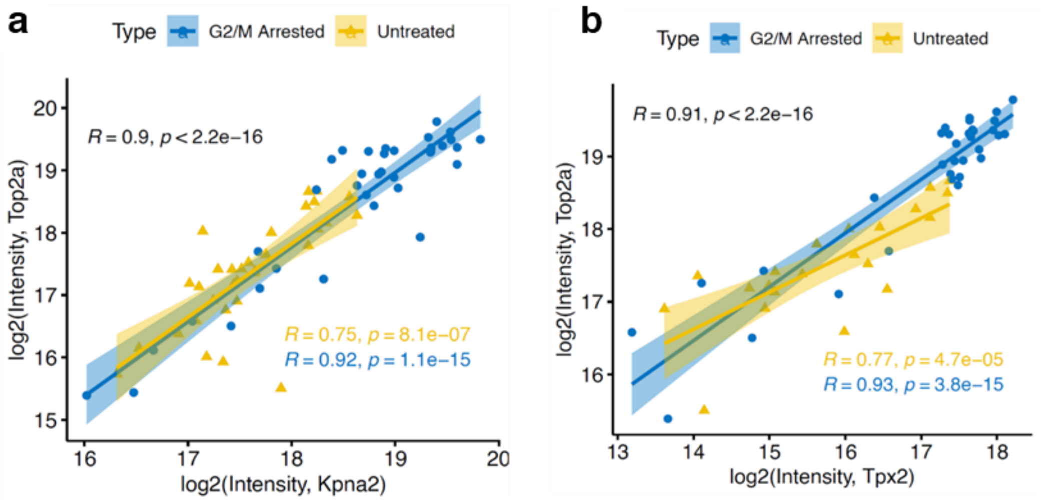


**Figure S26**: (**a**) Pearson correlations between KPNA2 and TOP2A using $log_2$(Intensity). Confidence intervals and data points are colored based on the cell group (G2/M arrested or untreated). Color also indicates Pearson correlation and p-value for each group, with black (upper left) representing the combined correlation of all cells in the plot. (**b**) Same as (**a**) except comparing TPX2 with TOP2A.

### Supplementary Results and Discussion

#### Concordance between scRNAseq and scProteomics in G2/M arrested cells

scProteomic covariant cluster 4 is notable in that several of the scProteomic fold-changes are reproduced in the scRNAseq results, demonstrating concordance between mRNA and protein (**Fig. 4b**). Three subclusters could be identified within cluster 4 based on protein covariation. These subclusters, though small in membership, provided statistically significant functional and pathway enrichment, with subcluster 4.1 showing clear enrichment for lysosomal processes and subcluster 4.2 being linked to hypoxia and metabolic pathways (**Fig. S20**). Proteins with close functional relationships (for example, prosaponin, progranulin, cathepsin A, and cathepsin D) exhibited strong correlations with each other (**Fig. S21a** and **S21b**). Finally, the largest clusters (5 and 6 from **Fig. 4b**) represented proteins with more modest differences in abundance for either condition.

While the scRNAseq data could cluster G2/M arrested and control C10 cells, we noted clustering of genes was less structured which we attributed to the stochastic nature of mRNA expression (**Fig. S11**). Despite limited overlap in differentially abundant genes and proteins (**Fig. S22**), many enriched GO terms were shared between scRNAseq and scProteomics (**Fig. S23**). For terms related to cell cycle and mitochondria, the combination of differentially expressed genes and proteins produced enrichments with greater statistical significance, suggesting the two modalities are more divergent in their shared identifications within these cases (**Fig. S23** and **Fig.S24**). Several protein clusters also produced enrichments with greater significance than what was found globally, and terms related to sphingolipid metabolism and ECM were exclusively found at the protein level (**Fig. S25**).

#### New insights into G2/M arrested cells from scProteomic data

One advantage of single-cell approaches is the identification of covarying protein or gene clusters in specific cell types or biological contexts[1,2]. In the context of our cell cycle arrest experiment, we identified several covarying clusters of proteins and genes including a cluster composed exclusively of canonical cell cycle proteins. Many of these proteins had exceptionally strong correlations, such as TOP2A, TPX2, and KPNA2 within G2/M arrested cells (**Fig. S26**). The strength of this correlation suggests the relative abundances of these proteins are tightly regulated. TPX2, a highly correlated protein within this cluster that is involved in spindle assembly, is known to be sequestered by importins-α/β with 1:1 stoichiometry through its nuclear localization sequence (NLS) before being released during G2 phase by HMMR (strongly correlated in this cluster as well) at the nuclear envelope [3,4]. Considering TOP2A contains a similar NLS[5], our data raises the possibility that TOP2A is regulated in a similar manner. Interestingly, many of the mitotic proteins described above did not show concordance with the scRNAseq data. One interpretation is that once the critical concentrations of these mitotic proteins are met, only lower levels of transcription are necessary to maintain proteins at that level. Furthermore, abundance (as measured by mRNA) of TOP2A, GTSE1, and CCNB1 has been shown to peak at G2 or in early mitosis[6,7] before rapid degradation at the protein level during mitotic exit[8]. Indeed, the G2/M arrested cells appear to be in the later stages of mitosis based on the mitotic protein cluster we identified[9]. Therefore, it can be inferred from our data that under cell cycle arrest the peak transcriptional states of many mitotic proteins are not continuously maintained.

We also observed a protein cluster containing ECM proteins (PCOLCE, FN1, COL1A1, COL3A1, and COL12A1), LNPK, and RIC8A decreasing in abundance in G2/M arrested cells. LNPK is known to be an endoplasmic reticulum(ER)-shaping protein, and reduction of LNPK facilitates the transition from tubular to sheet ER morphology during mitosis[10]. RIC8A is a key regulator during metaphase for mitotic spindle orientation, and reduction of Ric8a has been demonstrated to lead to prolonged mitosis and spindle defects[11]. As RIC8A was considerably less abundant in G2/M arrested cells, it may play a direct role in CDK1-inhibited mechanism of arrest. The remaining ECM proteins also shed light on cell morphological changes that occur during mitosis. It has been established that CDK1 regulates the remodeling of cell adhesion complexes during the cell cycle, and endogenous inactivation of CDK1 triggers this remodeling event[12,13]. Typically, this remodeling is thought to involve rapid recycling of cytoskeletal components. Our results demonstrated these ECM proteins are reduced at both the transcriptional and translational level during G2/M arrest, suggesting ECM degradation and transcriptional repression are also key to the adhesion complex remodeling process during mitosis. Hence, the nanoSPLITS-based single-cell multiomics not only reconstruct known mitotic processes but also identify new processes underlying biology.

### Supplementary Methods

#### nanoSPLITS buffer optimization

Using nuclease-free water (Thermo Fisher Scientific, cat# 4387936), several 10 mM Tris pH 8 test buffers were created containing 0.1% DDM and/or 1 x RNase inhibitor. 10 µL of each buffer was added to four wells within a 96 well plate. 20 C10 cells were then sorted into each well before snap freezing with liquid nitrogen. Immediately after thawing and centrifugation at 2,500 g, 5 µL from each well was transferred to a separate 96 well PCR plate containing 7.5 µL of 3' SMART-Seq CDS primer II before heating 70ºC for 3 min. 7.5 µL of RT mix (4 µL 5x ultra low first-strand buffer, 1 µL 48 µM SMART-Seq V4 oligonucleotide, 0.5 µL 40 units/µL RNase inhibitor, 2 µL SMARTScribe II reverse transcriptase) was then added before incubation at 42ºC for 90 min and 70ºC for 10 min. 30 µL of PCR master mix (25 µL SeqAmp PCR buffer, 1 µL PCR primer II A, 3 µL water, 1 µL SeqAmp DNA polymerase) was then added to each tube before performing 18 cycles of PCR (98ºC for 10 sec, 65ºC for 30 sec, 68ºC for 3 min). Isolation of cDNA was performed with Ampure XP beads with 80% ethanol washes. cDNA concentration and quality were determined with a Qubit fluorometer and Agilent fragment analyzer before next generation-sequencing, respectively.

The remaining 5 µL was retained and processed for label free proteomic analysis. Briefly, 5 µL of extraction buffer containing DTT and DDM was added to cell lysate to bring each sample to a final concentration of 1 mM DTT and 0.1% DDM before incubation at 60ºC for 1 h. 2 µL of 12 mM IAA was then added for a final concentration of 2 mM IAA before a 30 min incubation at 37ºC. 2 µL of 2.5 ng/µL Lys-C and 10 ng/µL of trypsin was added before incubation at 37ºC for 10 h. Enzymatic digestion was quenched by adding formic acid to a concentration of 1% before drying samples under vacuum. Peptides were reconstituted in 3 µL 5% acetonitrile 0.1% FA and transferred to a polypropylene microPOTS for proteomic analysis.

### Fluorescein nanoSPLITS split experiment and quantification

For the fluorescein-containing nanoSPLITS chip, 200 nL 0.01% 5,6-carboxyfluorescein solution containing 0.1% DDM was dispensed onto each well. For the PBS-containing chip, 250 nL PBS solution containing 0.1% DDM was dispensed onto each well. A slide cover was placed on both chips before wrapping them tightly in aluminum foil and placing them on ice to prevent evaporation until imaging. For imaging, both chips were placed on a chilled aluminum slide-holder and immediately imaged with brightfield light, followed by a Cy2 spectral filter using an AlphaImager FluorChemQ. Following imaging of the unsplit chips, two 2 $cm^2$ of 1/32" thick polyethylene foam was placed on one chip. The upper-chip was slowly lowered onto the bottom-chip, while carefully aligning the wells on both chips. Once the upper-chip was sitting on the separating foam, equal pressure was applied on the sides of the chip so that the droplets from both chips merged. Pressure was held for 15 seconds before releasing. The droplets were merged twice more following this process. The post-split chips were immediately placed back in the imager and final images were acquired. Quantification of droplet splitting was performed with the "Fiji" distribution of Image J (version 1.5). Briefly, Cy2 emission images were converted to grayscale. Regions of interest were selected (chip wells) and analyzed using the standard particle analysis in ImageJ. Each region of interest produced an average pixel intensity that was normalized by droplet area before using for quantification.

### Sparse partial least squares discriminant model (spls-DA) for identifying cell types in scProteomics data

Sparse partial least squares discriminant analysis (spls-DA) was utilized based on previously described algorithm[14]. From the scProteomic data, common contaminants were removed, and cells were filtered to have at least 3000 peptide identifications. Imputation was performed by sampling from a 3 standard deviation downshifted mean distribution. Three cell types were used to create different classes: alpha, beta, and delta (the remaining cells were considered as an alternative class).10-fold cross-validation was used to evaluate the model's predictive ability and balanced accuracy was used to assess sensitivity/specificity.
